# KDM2 proteins constrain transcription from CpG island gene promoters independently of their histone demethylase activity

**DOI:** 10.1101/561571

**Authors:** Anne H. Turberfield, Takashi Kondo, Manabu Nakayama, Yoko Koseki, Hamish W. King, Haruhiko Koseki, Robert J. Klose

## Abstract

CpG islands (CGI) are associated with the majority of mammalian gene promoters and function to recruit chromatin modifying enzymes. It has therefore been proposed that CGIs regulate gene expression through chromatin-based mechanisms, however in most cases this has not been directly tested. Here, we reveal that the histone H3 lysine 36 (H3K36) demethylase activity of the CGI-binding KDM2 proteins contributes only modestly to the H3K36me2-depleted state at CGI-associated gene promoters and is dispensable for normal gene expression. Instead, we discover that KDM2 proteins play a widespread and demethylase-independent role in constraining gene expression from CGI-associated gene promoters. We further show that KDM2 proteins shape RNA Polymerase II occupancy but not chromatin accessibility at CGI-associated promoters. Together this reveals a demethylase-independent role for KDM2 proteins in transcriptional repression and uncovers a new function for CGIs in constraining gene expression.

## INTRODUCTION

The functionality of complex multicellular organisms is underpinned by the creation of diverse cell types from a common genetic DNA blueprint. This is achieved during development by cells acquiring and maintaining cell type-specific gene expression programmes. At the most basic level, this relies on the control of RNA polymerase II (RNAPII)-mediated transcription by transcription factors (Spitz and Furlong 2012). However, it has also become clear that chromatin structure and its chemical modification can profoundly affect how transcription initiates from promoters and how gene expression is controlled (Kouzarides 2007; Li et al. 2007).

One such chemical modification of chromatin occurs on DNA where a methyl group is added to the 5 position of each cytosine in the context of CpG dinucleotides. CpG methylation is pervasive in mammalian genomes and is generally associated with transcriptional repression, particularly of repetitive and parasitic DNA elements (Klose and Bird 2006; Schübeler 2015). However, short CpG-rich regions of the genome, called CpG islands (CGIs), remain free of DNA methylation and are associated with the majority of mammalian gene promoters (Saxonov et al. 2006; Illingworth and Bird 2009). CGIs have been proposed to regulate gene expression (Blackledge and Klose 2011; Deaton and Bird 2011) through a family of ZF-CxxC DNA binding domain-containing proteins that recognise non-methylated DNA and occupy CGIs (Voo et al. 2000; Lee et al. 2001; Blackledge et al. 2010; Thomson et al. 2010). Interestingly, most ZF-CxxC domain-containing proteins possess histone modifying activities or are part of large chromatin modifying complexes, suggesting that these factors regulate gene expression through chromatin (Long et al. 2013a). However, in most cases the contribution of chromatin-based mechanisms to CGI-dependent gene regulation remains untested.

The ZF-CxxC domain-containing protein lysine-specific demethylase 2A (KDM2A) and its paralogue KDM2B bind to CGIs (Blackledge et al. 2010; Farcas et al. 2012; He et al. 2013; Wu et al. 2013). KDM2 proteins encode a JmjC domain that catalyses the removal of H3K36 mono- and dimethylation (H3K36me1/2) (Tsukada et al. 2006; Fang et al. 2007; He et al. 2008; Cheng et al. 2014). H3K36me1/2 are broadly distributed throughout the mammalian genome (Peters et al. 2003; Robin et al. 2007; Schotta et al. 2008) and H3K36me2 has been proposed to counteract transcription initiation. For example, in yeast H3K36me2 inhibits inappropriate initiation of transcription from cryptic promoters in genes (Carrozza et al. 2005; Joshi and Struhl 2005; Keogh et al. 2005; Li et al. 2009a; McDaniel and Strahl 2017), and this function may also be conserved in mammals (Xie et al. 2011; Carvalho et al. 2013). Given the seemingly widespread and indiscriminate deposition of H3K36me1/2 in mammalian genomes and its association with transcriptional repression, the discovery that KDM2 proteins localise specifically to CGIs has led to the suggestion that the removal of H3K36me2 at these sites may contribute to a widespread and transcriptionally permissive chromatin state at gene promoters (Blackledge and Klose 2011; Deaton and Bird 2011). Depletion of KDM2A was shown to cause an increase in H3K36me2 at a number of CGI-associated gene promoters, suggesting that KDM2A plays an active role in H3K36me2 removal (Blackledge et al. 2010). However, the histone demethylase activity of KDM2 proteins has also been linked to gene repression, including of genes that have roles in cell proliferation, differentiation and senescence (Frescas et al. 2007; He et al. 2008; Tzatsos et al. 2009; Tanaka et al. 2010; Du et al. 2013; Yu et al. 2016). Therefore, the role that KDM2 proteins play in regulating H3K36me1/2 and the effect that this has on CGI-associated gene transcription remain unclear.

KDM2 proteins may also regulate transcription through mechanisms that do not rely on their demethylase activity. KDM2A has been implicated in the formation of pericentromeric heterochromatin (Borgel et al. 2017), while KDM2B physically associates with polycomb repressive complex 1 (PRC1) and is required for the formation of repressive polycomb chromatin domains at a subset of CGI-associated gene promoters (Farcas et al. 2012; He et al. 2013; Wu et al. 2013; Blackledge et al. 2014). The possibility that KDM2 proteins have demethylase-independent activity is supported by the observation that the *Kdm2a* and *Kdm2b* genes encode internal transcription start sites (TSS) downstream of their JmjC domain. Transcription initiating from these alternative promoters gives rise to short forms of KDM2A and KDM2B (KDM2A/B-SF, Figure 1A) that lack the JmjC domain and therefore cannot act as histone demethylases (Tanaka et al. 2010; Long et al. 2013a). Importantly, however, they retain their ZF-CxxC domain and CGI binding activity. The function of KDM2A/B-SF proteins remains poorly defined, but there is evidence that the KDM2B-SF is sufficient to recruit PRC1 to chromatin (Blackledge et al. 2014). Following the depletion of either KDM2A or KDM2B, alterations in gene expression have been reported (Blackledge et al. 2010; Farcas et al. 2012; He et al. 2013; Blackledge et al. 2014; Boulard et al. 2015). However, whether these two closely related paralogues function cooperatively to regulate gene expression is unknown and, like many chromatin modifying enzymes, it remains largely untested whether they rely on their enzymatic activity for gene regulation. Perhaps more fundamentally, whether the KDM2 proteins function primarily to potentiate or repress gene transcription has not been examined at the genome-scale and remains a major conceptual barrier in understanding how CGIs, which are associated with most vertebrate gene promoters, control gene transcription and expression.

**Figure 1.**
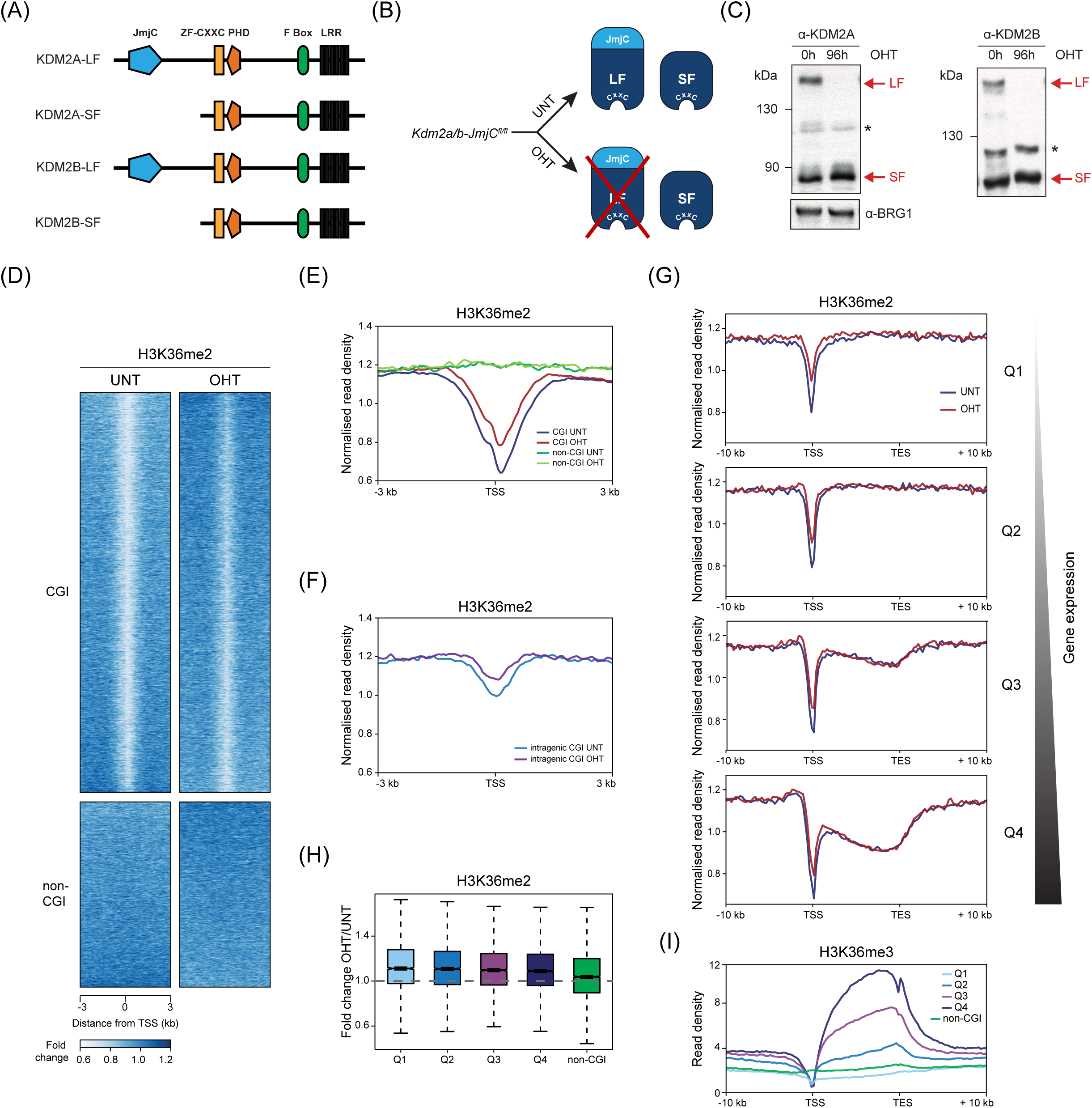
KDM2 proteins contribute modestly to the H3K36me2-depleted state at CGI-associated gene promoters. (A) A schematic illustrating protein domain architecture for KDM2A/B long (LF) and short isoforms (SF). (B) A schematic of the *Kdm2a/b-JmjC*^*fl/fl*^ system in which addition of tamoxifen (OHT) leads to removal of KDM2 long isoforms. (C) Western blot analysis for KDM2A and KDM2B in K*dm2a/b-JmjC*^*fl/fl*^ mESCs before (UNT) and after 96 hours of tamoxifen (OHT) treatment. BRG1 is shown as a loading control for both blots. Asterisks indicate non-specific bands. (D) Heatmaps of H3K36me2 enrichment (ChIP-seq) in K*dm2a/b-JmjC*^*fl/fl*^ mESCs before (UNT) and after addition of tamoxifen (OHT), for CGI-associated (n=14106) and non-CGI-associated (n=6527) gene promoters. H3K36me2 signal was normalised to H3 ChIP-seq to control for any alterations in nucleosome density. (E) A metaplot of normalised H3K36me2 ChIP-seq signal at CGI-associated or non-CGI-associated gene promoters in K*dm2a/b-JmjC*^*fl/fl*^ mESCs, before (UNT) and after tamoxifen treatment (OHT). (F) A metaplot of normalised H3K36me2 ChIP-seq signal at intragenic CGIs in K*dm2a/b-JmjC*^*fl/fl*^ mESCs, before (UNT) and after tamoxifen treatment (OHT). (G) Metaplots showing normalised H3K36me2 ChIP-seq signal throughout the gene body before (UNT) and after tamoxifen treatment (OHT), for CGI-associated genes separated into quartiles according to their expression level in K*dm2a/b-JmjC*^*fl/fl*^ mESCs (Q1 < Q2 < Q3 < Q4). Genes were scaled to the same length and aligned at their TSS and TES. (H) A boxplot showing fold change in normalised H3K36me2 ChIP-seq signal following tamoxifen treatment, for the CGI-associated gene quartiles shown in (G) and for non-CGI-associated genes. (I) A metaplot showing H3K36me3 enrichment throughout the gene body for the gene sets shown in (G) (Brookes et al. 2012).

To address these fundamental questions, here we have exploited systematic conditional genetic ablation strategies and detailed genome-wide analysis to dissect how KDM2 proteins regulate H3K36me2 and gene expression in mouse embryonic stem cells (mESCs). Remarkably, we discover that KDM2 proteins contribute only modestly to the H3K36me2-depleted state at CGI-associated gene promoters and the demethylase activity of KDM2 proteins is largely dispensable for normal gene expression. In contrast, surgical removal of the KDM2 ZF-CxxC domains, which liberates KDM2 proteins from CGIs, revealed a widespread increase in gene expression. This was not limited to the function of KDM2B in polycomb-mediated gene repression, but instead occurred broadly across CGI-associated genes, revealing an unexpectedly widespread role for KDM2 proteins in constraining gene expression. KDM2B plays the predominant role in gene repression, while KDM2A appears to cooperate with KDM2B to counteract expression at a subset of genes. KDM2-dependent effects on gene expression do not manifest through altered DNA accessibility at CGIs, but instead appear to regulate RNAPII occupancy at gene promoters. Therefore, we define a new demethylase-independent role for KDM2A/B in transcriptional repression, uncovering a new logic whereby CGIs appear, unexpectedly, to constrain gene expression.

## RESULTS

### KDM2 proteins contribute modestly to the H3K36me2-depleted state at CGI-associated gene promoters

KDM2A and KDM2B both catalyse H3K36me2 demethylation via their JmjC domain (Tsukada et al. 2006; He et al. 2008) and localise to CGIs via their ZF-CxxC domain (Figure 1A) (Blackledge et al. 2010; Farcas et al. 2012; He et al. 2013; Wu et al. 2013). However, whether KDM2 proteins regulate H3K36me2 at CGI-associated gene promoters throughout the genome has not been examined. Therefore, we generated a mESC system in which loxP sites were inserted into the *Kdm2a* and *Kdm2b* genes flanking exons that encode the JmjC domain (*Kdm2a/b-JmjC*^*fl/fl*^, Supplementary Figure 1A) and which also expresses a tamoxifen-inducible form of CRE recombinase. Addition of tamoxifen triggers deletion of the JmjC domain-containing exons, removing the long forms of KDM2A and KDM2B (KDM2-LFs) and their associated demethylase activity (Figure 1B,C, Supplementary Figure 1B). Importantly, KDM2-SFs, which are expressed from downstream promoters (Supplementary Figure 1A), were unaffected by removal of the KDM2-LFs (Figure 1 B,C). We first investigated the contribution of KDM2-LFs to global H3K36 methylation levels by western blot, and observed only minor changes following removal of KDM2-LFs (Supplementary Figure 1C,D). Next, we examined the genome-wide distribution of H3K36me2 using chromatin immunoprecipitation followed by massively-parallel sequencing (ChIP-seq). This confirmed a local depletion of H3K36me2 at CGI-associated gene promoters (Figure 1D,E) (Blackledge et al. 2010; Blackledge and Klose 2011; Deaton and Bird 2011). H3K36me2 depletion was not detected at non-CGI gene promoters, demonstrating that this is a CGI-associated chromatin feature. Following removal of the KDM2-LFs by tamoxifen treatment, there was a modest increase in H3K36me2 at the TSS of CGI-associated gene promoters (Figure 1D,E). This demonstrates that KDM2A/B contribute to the H3K36me2-depleted state at CGIs, in agreement with single-gene studies examining the KDM2A- or KDM2B-depleted state (Blackledge et al. 2010). Interestingly, intragenic CGIs were also depleted of H3K36me2 (Figure 1F), although this depletion was on average less pronounced than at CGI promoters, likely due to their lower average CpG density and size (Supplementary Figure 1E). Importantly, removal of the KDM2-LFs resulted in an increase in H3K36me2 at intragenic CGIs indicating that their H3K36me2 depleted state is also shaped by KDM2A/B.

Our ChIP-seq analysis revealed that KDM2-LFs contribute to the H3K36me2-depleted state at CGI-associated promoters. However, we were curious whether the effects on H3K36me2 were uniformly distributed or dependent on other features of gene promoters, such as transcriptional activity. Therefore, we separated genes based on expression level (Supplementary Figure 1F) and examined H3K36me2 at genes and surrounding regions. This revealed that CGI-associated TSSs were depleted of H3K36me2 irrespective of expression level (Figure 1G). Chromatin surrounding CGI-associated TSSs was blanketed by H3K36me2, consistent with this modification being pervasive in mammalian genomes. The increase in H3K36me2 at the TSS following KDM2-LF removal was similar across all expression levels (Figure 1H), consistent with the transcription-independent targeting of KDM2 proteins to CGI promoters via their ZF-CxxC domains. Interestingly, highly transcribed genes were also depleted of H3K36me2 in their gene body (Figure 1G). However, this was independent of KDM2 demethylase activity and instead correlated with co-transcriptional conversion of H3K36me2 to H3K36me3 (Figure 1I) (Bannister et al. 2005; Pokholok et al. 2005; Barski et al. 2007; Bell et al. 2007; Mikkelsen et al. 2007; Weiner et al. 2015). Together, these observations reveal that KDM2 proteins remove H3K36me2 from CGIs, but unexpectedly their contribution to the depletion of H3K36me2 at these sites is modest. This suggests that CGIs could be inherently refractory to H3K36me2 or that additional H3K36 demethylases may also function at these regions (see discussion).

### KDM2 demethylase activity contributes minimally to gene regulation

Depletion of H3K36me2 at CGI-associated gene promoters has been proposed to contribute to the generation of a transcriptionally permissive chromatin state (Blackledge et al. 2010; Blackledge and Klose 2011; Deaton and Bird 2011). Although KDM2 proteins appear to contribute only modestly to the H3K36me2-depleted state at CGIs (Figure 1), we were curious whether this effect was nevertheless required to sustain normal chromatin accessibility and transcription from CGI-associated gene promoters. To address these questions, we first performed calibrated ATAC-seq (cATAC-seq) to measure chromatin DNA accessibility before and after removal of KDM2-LFs. This demonstrated that CGI promoters remained accessible, despite the observed increases in H3K36me2 (Figure 2A). To examine gene expression, we performed calibrated nuclear RNA sequencing (cnRNA-seq). This revealed that the expression of the vast majority of genes did not change following removal KDM2 demethylase activity, with only a small number of genes being modestly perturbed (Figure 2B). Furthermore, there was a poor correlation between gene expression changes and the effects on H3K36me2 at gene promoters (Figure 2C,D). This minimal perturbation to gene expression and chromatin accessibility following loss of KDM2-LFs indicates that histone demethylase activity of KDM2 proteins is largely dispensable for normal CGI-associated promoter activity.

**Figure 2.**
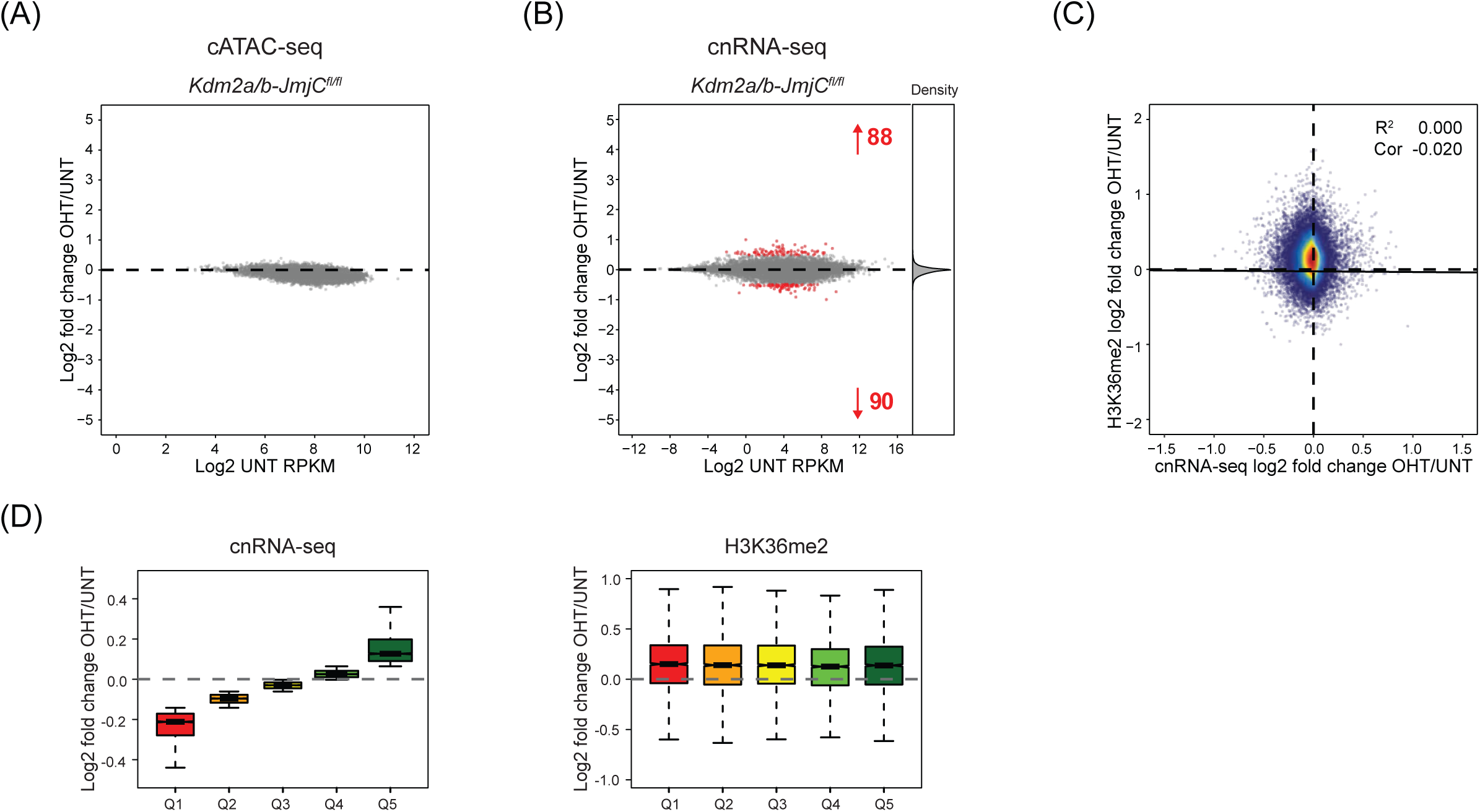
KDM2 demethylase activity contributes minimally to gene regulation. (A) An MA-plot showing log2 fold change in the accessibility (cATAC-seq) of CGI-associated gene promoters in K*dm2a/b-JmjC*^*fl/fl*^ mESCs following tamoxifen treatment. No promoters significantly changed in accessibility (p-adj < 0.05 and > 1.4-fold). (B) An MA-plot showing log2 fold change in gene expression (cnRNA-seq) in K*dm2a/b-JmjC*^*fl/fl*^ mESCs following tamoxifen treatment. The number of genes with significantly increased or decreased expression (p-adj < 0.05 and > 1.4-fold) is shown in red and density of gene expression changes is shown on the right. (C) A scatter plot comparing the log2 fold change in gene expression (cnRNA-seq) with the log2 fold change in normalised H3K36me2 ChIP-seq signal for CGI-associated genes following tamoxifen treatment of *Kdm2a/b-JmjC*^*fl/fl*^ mESCs. The solid line shows the linear regression, and the coefficient of determination (R^2^) and Spearman correlation coefficient (Cor) are annotated. (D) Boxplots showing the log2 fold change in cnRNA-seq signal (left) and normalised H3K36me2 ChIP-seq signal (right) for CGI-associated genes grouped into quintiles based on their change in expression following tamoxifen treatment of *Kdm2a/b-JmjC*^*fl/fl*^ mESCs.

### KDM2 proteins play a widespread role in gene repression

Given that gene expression was largely unaffected when KDM2 demethylase activity was removed, we wondered whether demethylase-independent activities of KDM2 proteins may play a more prominent role in gene regulation. KDM2A and KDM2B encode multiple isoforms, each of which contain the ZF-CxxC DNA binding domain. Therefore, to remove all CGI-targeted KDM2 proteins, we developed a conditional mESC system in which the exon encoding the ZF-CxxC domain is flanked by loxP sites in both *Kdm2a* and *Kdm2b* genes (*Kdm2a/b-CXXC*^*fl/fl*^), and which expresses tamoxifen-inducible Cre recombinase (Supplementary Figure 3A). Following addition of tamoxifen the ZF-CxxC-encoding exons are excised, producing KDM2A and KDM2B proteins that now lack the ZF-CxxC domain (Figure 3A). The effectiveness of this approach was evident from the loss of the ZF-CxxC domain and an increased mobility of the KDM2 proteins in western blot analysis (Figure 3B) and from loss of binding to CGIs in ChIP analysis (Supplementary Figure 3B).

**Figure 3.**
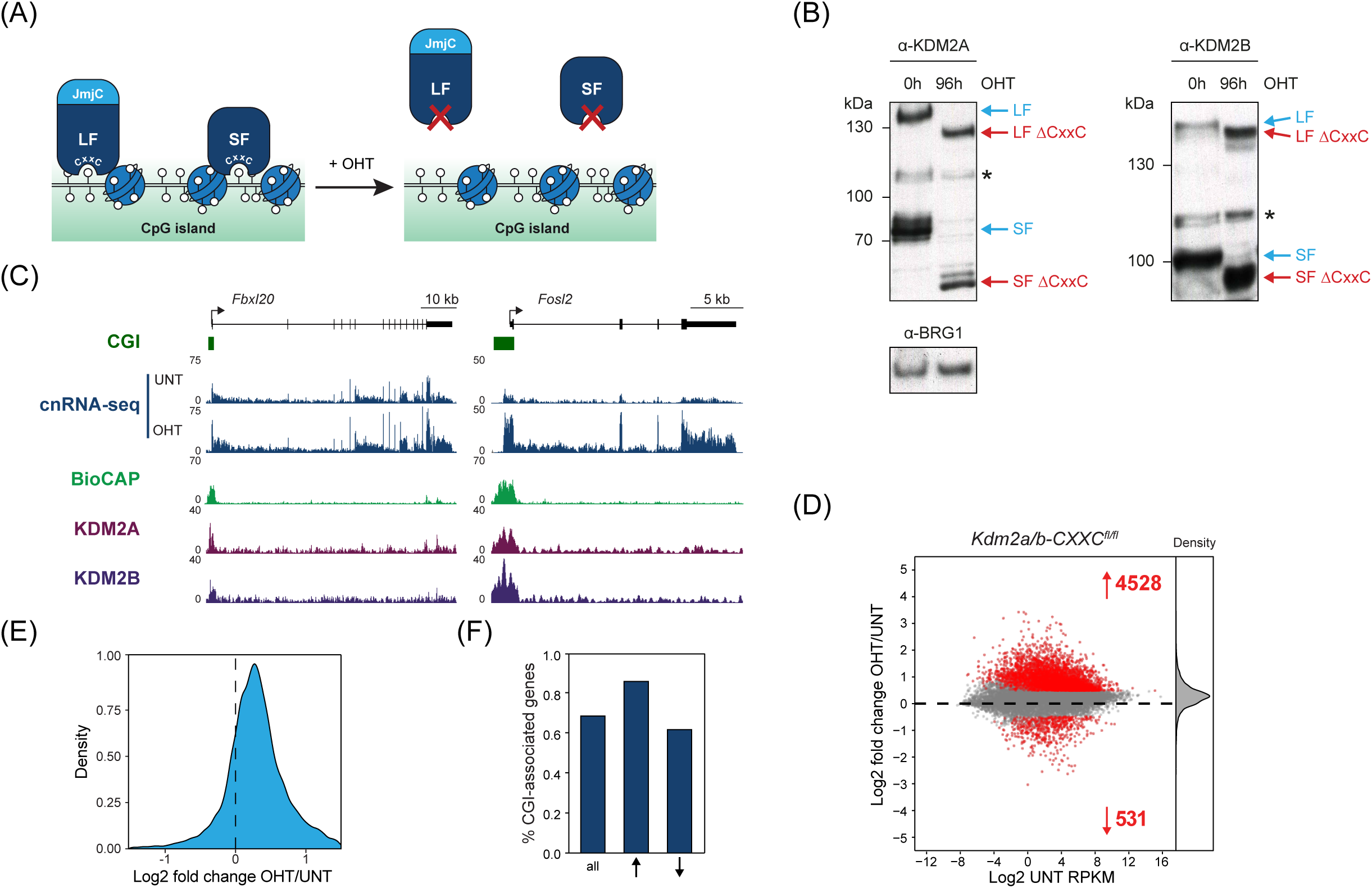
KDM2 proteins mediate widespread gene repression. (A) A schematic of the *Kdm2a/b-CXXC*^*fl/fl*^ system in which addition of tamoxifen (OHT) leads to the generation of KDM2 proteins that lack the ZF-CxxC domain and therefore are unable to bind to chromatin. (B) Western blot analysis for KDM2A and KDM2B in *Kdm2a/b-CXXC*^*fl/fl*^ mESCs before (UNT) and after 96 hours tamoxifen (OHT) treatment. BRG1 is shown as a loading control for both blots. Asterisks indicate non-specific bands. (C) Genomic snapshots showing gene expression (cnRNA-seq) before (UNT) and after tamoxifen (OHT) treatment of *Kdm2a/b-CXXCfl/fl* mESCs, for representative genes that moderately (*Fbxl20*, left) or more dramatically increased in expression (*Fosl2*, right). BioCAP and KDM2A and KDM2B ChIP-seq signal are shown for reference (Farcas et al. 2012; Long et al. 2013b; Blackledge et al. 2014) (D) An MA-plot showing log2 fold change in gene expression (cnRNA-seq) in K*dm2a/b-CXXC*^*fl/fl*^ mESCs following tamoxifen treatment. The number of genes with significantly increased or decreased expression (p-adj < 0.05 and > 1.4-fold) are shown in red and density of gene expression changes is shown on the right. (E) A density plot showing the distribution of the log2 fold change in gene expression following tamoxifen treatment of *Kdm2a/b-CXXC*^*fl/fl*^ mESCs, for all genes. (F) A bar graph showing the proportion of genes that have a CGI promoter, for all genes and genes that significantly increased or decreased in expression following tamoxifen treatment of *Kdm2a/b-CXXC*^*fl/fl*^ mESCs.

To examine whether this loss of CGI binding had an effect on gene expression, we carried out cnRNA-seq and compared gene expression between untreated and tamoxifen treated cells. This revealed that KDM2 protein removal resulted in more than a fifth of all genes showing significantly increased expression (Figure 3C,D). Owing to the quantitative nature of cnRNA-seq it was also apparent that KDM2 protein removal led to a more general increase in gene expression, even amongst genes that were not considered significantly changed by statistical analysis (Figure 3E). We validated these widespread effects using highly sensitive and quantitative digital droplet PCR analysis (Supplementary Figure 3C). Importantly, our capacity to uncover this broad increase in gene expression was only possible due to the use of calibrated nuclear RNA-seq (cnRNA-seq) as conventional normalisation based on total read count fails to uncover this pervasive alteration in gene expression (Supplementary Figure 3D). When we examined in more detail the transcripts with significantly increased expression these were enriched for CGI-associated genes (Figure 3F), consistent with these effects being a direct result of KDM2 protein removal as opposed to a global perturbation of some core transcriptional component. In contrast, significantly downregulated genes were less numerous and not enriched for CGI-associated genes, suggesting they may correspond to secondary effects. Together, these observations establish an unexpected and widespread role for KDM2 proteins in suppressing expression from CGI-associated gene promoters.

### Elevated gene expression following KDM2 protein removal is not simply a consequence of polycomb target gene reactivation

We have previously shown that KDM2B plays an important role in recruiting the PRC1 complex to CGI-associated gene promoters in mESCs. It does so by interacting with the PRC1 complex via the specialised adaptor protein PCGF1 which links KDM2B to RING1B, the catalytic core of PRC1 (Farcas et al. 2012; He et al. 2013; Wu et al. 2013; Blackledge et al. 2014). When we examined the genes that increased in gene expression following KDM2 protein removal they had stereotypical CGI-associated features (Figure 4A), but were also enriched for KDM2B, RING1B and SUZ12. This raised the possibility that the observed effects on gene expression following KDM2 protein removal simply resulted from loss of KDM2B-dependent targeting and gene repression by the PRC1 complex. To investigate this possibility, we compared the gene expression changes following KDM2 protein removal with those following conditional removal of PCGF1 (Fursova 2019). This revealed that removal of PCGF1 caused de-repression of more than four times fewer genes than removal of KDM2 proteins (Figure 4B). Furthermore, genes that significantly increased in expression following PCGF1 removal were more strongly enriched for polycomb target genes than those that significantly increased following KDM2 removal (Figure 4C). Genes showing increased expression following PCGF1 removal were a subset of those showing increased expression following KDM2 protein removal (Figure 4D), and there was only a moderate positive correlation between the gene expression changes in these lines (Supplementary Figure 4A). These observations indicate that a small proportion of the gene de-repression events in cells where KDM2 proteins are removed are related to the activity of the KDM2B-PRC1 complex.

**Figure 4.**
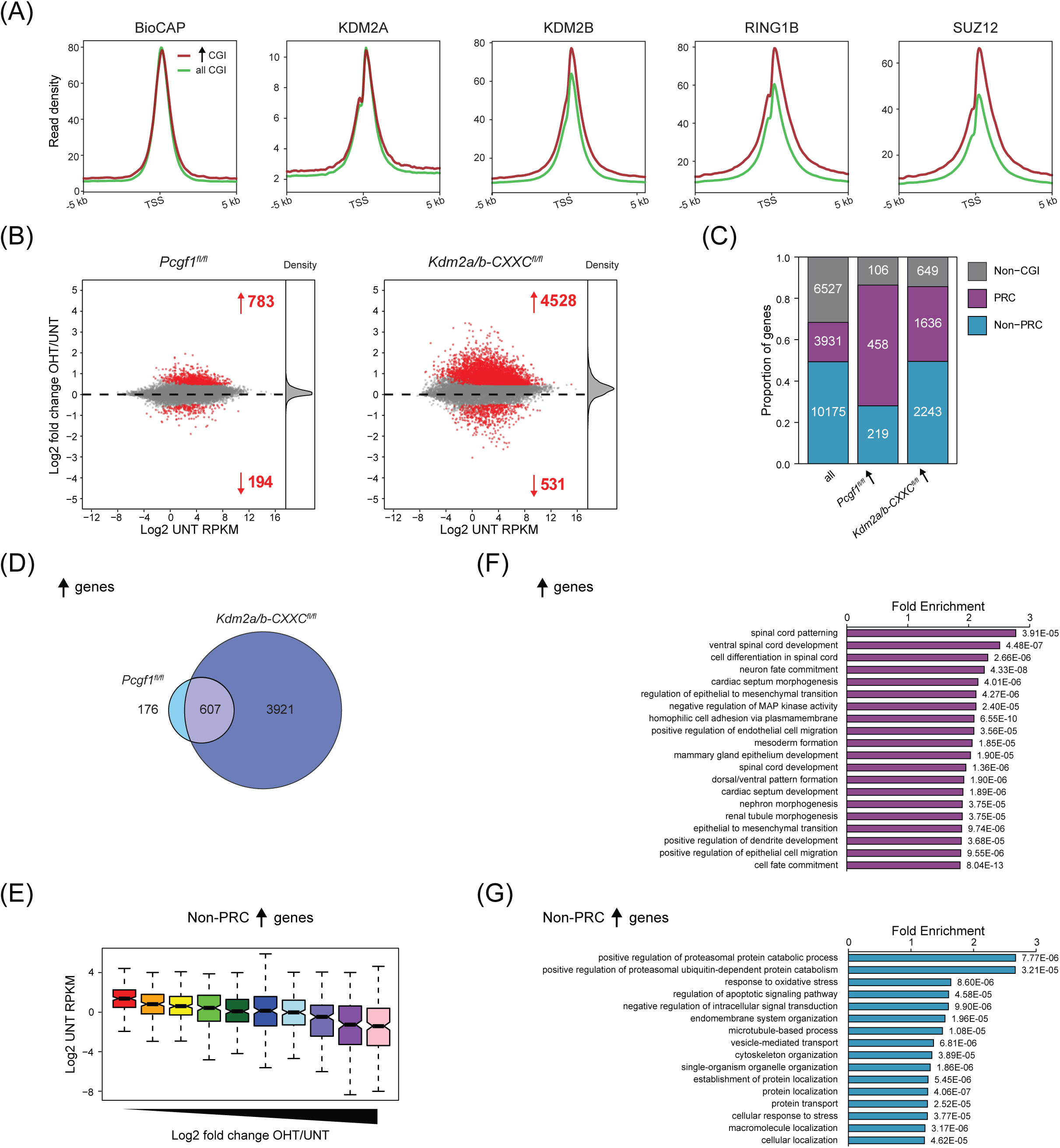
KDM2-mediated repression is not limited to polycomb target genes. (A) Metaplots showing enrichment of BioCAP signal and KDM2A, KDM2B, RING1B and SUZ12 ChIP-seq signal at the TSS of all CGI-associated genes (n=14106, green) and of the subset of these genes that significantly increased in expression following tamoxifen treatment of *Kdm2a/b-CXXC*^*fl/fl*^ mESCs (n=3879, red) (Farcas et al. 2012; Long et al. 2013b; Blackledge et al. 2014). (B) Left: an MA-plot showing log2 fold change in gene expression (cnRNA-seq) in *Pcgf1*^*fl/fl*^ mESCs following tamoxifen treatment to induce PCGF1 knockout (Fursova 2019). The number of genes with significantly increased or decreased expression (p-adj < 0.05 and > 1.4-fold) is shown in red and density of gene expression changes is shown on the right. Right: as (B) but for *Kdm2a/b-CXXC*^*fl/fl*^ mESCs, shown for comparison. (C) A bar graph comparing the distribution of genes into three classes – non-CGI, polycomb (PRC) occupied and non-PRC occupied – for all genes and for genes that that significantly increased in expression following tamoxifen treatment of *Pcgf1*^*fl/*fl^ or *Kdm2a/b-CXXC*^*fl/fl*^ mESCs. Non-CGI genes are genes that lack a CGI at their promoter. Non-PRC-occupied genes have a CGI promoter that is not bound by polycomb complexes, while PRC-occupied genes have a CGI promoter that is bound by polycomb complexes. (D) A Venn diagram showing the overlap between genes that significantly increased in expression following tamoxifen treatment of *Pcgf1*^*fl/*fl^ and *Kdm2a/b-CXXC*^*fl/fl*^ mESCs. (E) A box plot showing the starting expression level (log2 UNT RPKM) for genes grouped into deciles based on their log2 fold change in expression following tamoxifen treatment of *Kdm2a/b-CXXC*^*fl/fl*^ mESCs. (F) Gene ontology analysis of genes that significantly increased in expression following tamoxifen treatment of *Kdm2a/b-CXXC*^*fl/fl*^ mESCs. (G) As (H), but for the subset of significantly increasing genes that were not classified as polycomb target genes.

Building on this important observation, we examined in more detail the gene expression changes that manifest from KDM2 protein removal from CGIs. From this it was evident that genes with low starting expression level more strongly increased in expression, including non-polycomb target genes (Figure 4E). Gene ontology analysis revealed that genes that significantly increased in expression were enriched for a variety of developmental terms (Figure 4F), consistent with some of the effects being related to the polycomb system, but also a variety of basic cellular processes that are unrelated (Figure 4G). This reflects the generalised increase in gene expression that occurs following KDM2 removal. Together, these observations reveal that KDM2 proteins play a widespread role in gene repression from CGI-associated gene promoters and do so largely through mechanisms that are independent of the polycomb repressive system.

### KDM2B plays the predominant role in gene repression

Loss of both KDM2A and KDM2B from CGI chromatin simultaneously resulted in widespread increases in gene expression (Figure 3). However, it was unclear from these experiments whether KDM2A, KDM2B or both contribute to gene repression. To examine this question, we developed a conditional mESC system in which we could remove KDM2A alone by tamoxifen-induced deletion of its ZF-CxxC domain (*Kdm2a-CXXC*^*fl/fl*^, Figure 5A, Supplementary Figure 5A and see Supplementary Figure 3A). cnRNA-seq revealed that removal of KDM2A led to virtually no changes in gene expression (Figure 5B), indicating that KDM2A alone is not required to maintain normal gene expression in mESCs. We next investigated the contribution of KDM2B to the regulation of gene expression, performing cnRNA-seq using a *Kdm2b-CXXC*^*fl/fl*^ mESC line (Blackledge et al. 2014)(Figure 5A, Supplementary Figure 5B). cnRNA-seq revealed that removal of KDM2B alone was sufficient to cause increases in the expression of thousands of genes (Figure 5B) and, unlike KDM2A removal, largely recapitulated the widespread increases in gene expression that occurred following KDM2A/B removal (Figure 5C). Genes that significantly increased in expression following KDM2B removal were enriched for CGI-associated genes (Figure 5D). Furthermore, gene ontology analysis revealed that these significantly increasing genes were enriched for a variety of developmental terms characteristic of polycomb target genes (Figure 5E) but also terms relating to basic cellular processes (Figure 5F). These observations suggest that removal of KDM2B alone, like removal of KDM2A/B together, leads to widespread increases in the expression of CGI-associated genes.

**Figure 5.**
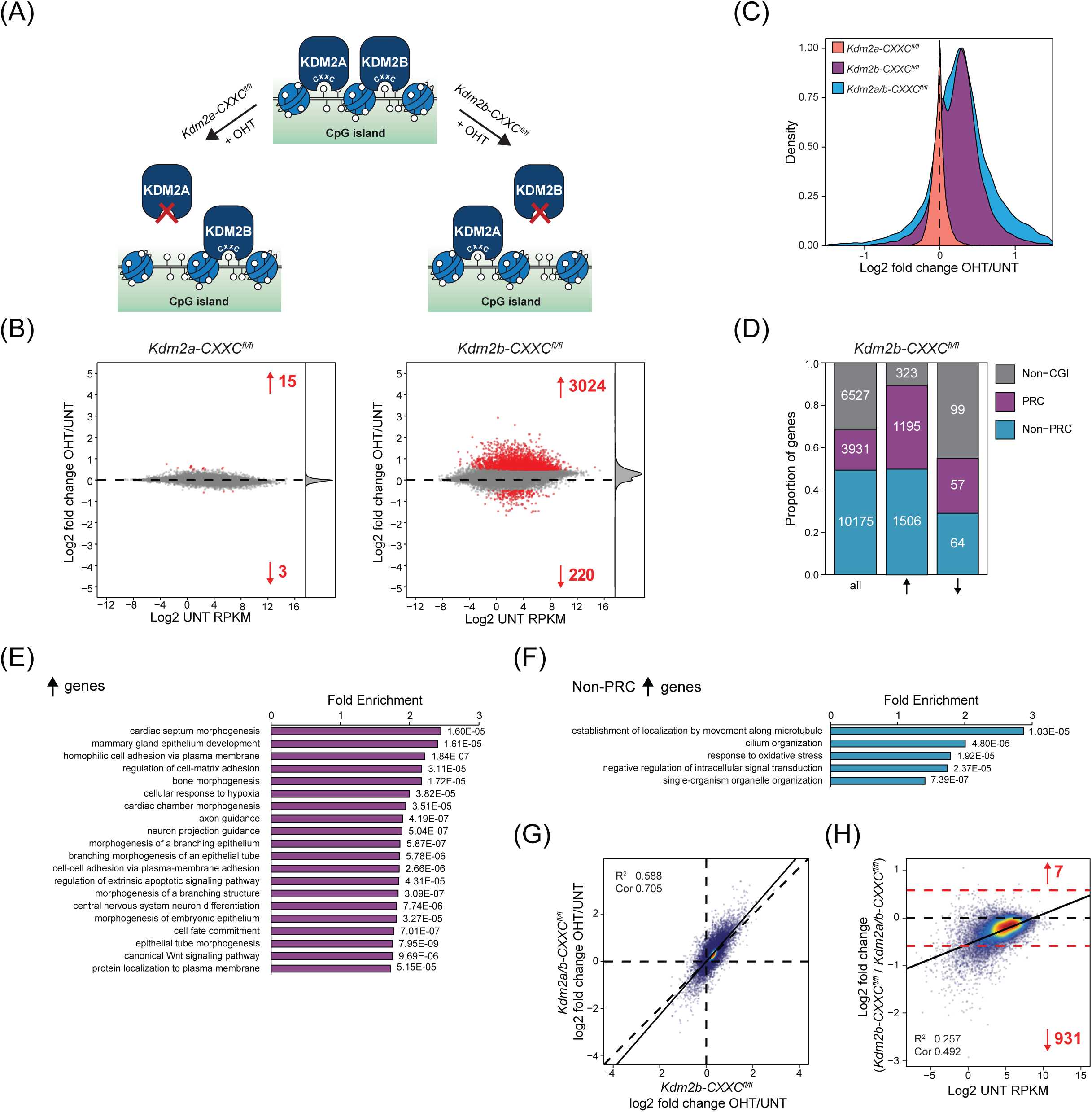
KDM2B plays the predominant role in gene repression. (A) A schematic of the *Kdm2a-CXXC*^*fl/fl*^ and *Kdm2b-CXXC*^*fl/fl*^ systems in which addition of tamoxifen (OHT) leads to the generation of KDM2A or KDM2B proteins that lack the ZF-CxxC domain, respectively, and therefore are unable to bind to chromatin. (B) MA-plots showing log2 fold change in gene expression (cnRNA-seq) in *Kdm2a-CXXC*^*fl/fl*^ (left) or *Kdm2b-CXXC*^*fl/fl*^ mESCs (right) following tamoxifen treatment. The number of genes with significantly increased or decreased expression (p-adj < 0.05 and > 1.4-fold) are shown in red and density of gene expression changes is shown on the right. (C) A density plot showing the distribution of the log2 fold change in gene expression following tamoxifen treatment of *Kdm2a-CXXC*^*fl/fl*^, *Kdm2b-CXXC*^*fl/fl*^ or *Kdm2a/b-CXXC*^*fl/fl*^ mESCs, for all genes. (D) A bar graph illustrating the distribution of genes between three gene classes (Non-CGI, Non-PRC, PRC) described in Figure 4C, for all genes and for genes that significantly increased or decreased in expression following tamoxifen treatment of *Kdm2b-CXXC*^*fl/fl*^ mESCs. (E) Gene ontology analysis of genes that significantly increased in expression following tamoxifen treatment of *Kdm2b-CXXC*^*fl/fl*^ mESCs. (F) As (E), but for the subset of significantly increasing genes that were not classified as polycomb target genes. (G) A scatter plot comparing the log2 fold change in gene expression (cnRNA-seq) following tamoxifen treatment of *Kdm2b-CXXC*^*fl/fl*^ and *Kdm2a/b-CXXC* ^*fl/fl*^ mESCs. The solid line shows the linear regression, and the coefficient of determination (R^2^) and Spearman correlation coefficient (Cor) are annotated. (H) A scatter plot of the log2 fold change in gene expression (cnRNA-seq) following tamoxifen treatment of *Kdm2a/b-CXXC* ^*fl/fl*^ mESCs for genes that significantly increased in expression, plotted against the ratio of the log2 fold change in gene expression following tamoxifen treatment of *Kdm2b-CXXC* ^*fl/fl*^ versus *Kdm2a/b-CXXC* ^*fl/fl*^ mESCs. A 1.5-fold threshold (red dotted lines) was used to define genes which were differentially regulated between the two datasets, and the number of genes with more than 1.5-fold increased or decreased expression is shown in red. The solid line shows the linear regression, and the coefficient of determination (R^2^) and Spearman correlation coefficient (Cor) are annotated.

A comparison of the gene expression changes following KDM2B removal alone and KDM2A/B removal together revealed good overall correlation (Figure 5G), indicating that the gene expression changes following KDM2B removal largely recapitulated those following removal of KDM2A/B together. However, a more detailed analysis revealed 931 significantly increasing genes that less strongly increased in expression following loss of KDM2B compared to KDM2A/B together, and this set was enriched for genes with low expression level (Figure 5H). This suggests that KDM2A plays a role in restricting the expression of these genes following KDM2B removal. Together our findings demonstrate that KDM2B plays the predominant role in repressing gene expression, while KDM2A may cooperate with KDM2B to counteract expression at a subset of genes.

### KDM2 proteins regulate polymerase occupancy but not chromatin accessibility at CGIs

Gene regulatory elements and gene promoters are characterised by elevated chromatin accessibility (Boyle et al. 2008; Song et al. 2011; Thurman et al. 2012), and this is thought to play an important role in regulating gene expression. Accessibility at CGI-associated gene promoters broadly correlates with transcriptional output, with the promoters of highly transcribed CGI-associated genes being more highly accessible (King et al. 2018). Therefore we wondered whether the increases in gene expression following removal of KDM2 proteins resulted from increases in the accessibility at CGI-associated gene promoters in the absence of KDM2A/B. To test this we carried out cATAC-seq following removal of KDM2 proteins. Importantly, we did not observe any significant change in the accessibility of CGI-associated gene promoters (Figure 6A, Supplementary Figure 6A), indicating that expression changes must manifest through effects on transcription that are independent of chromatin accessibility.

**Figure 6.**
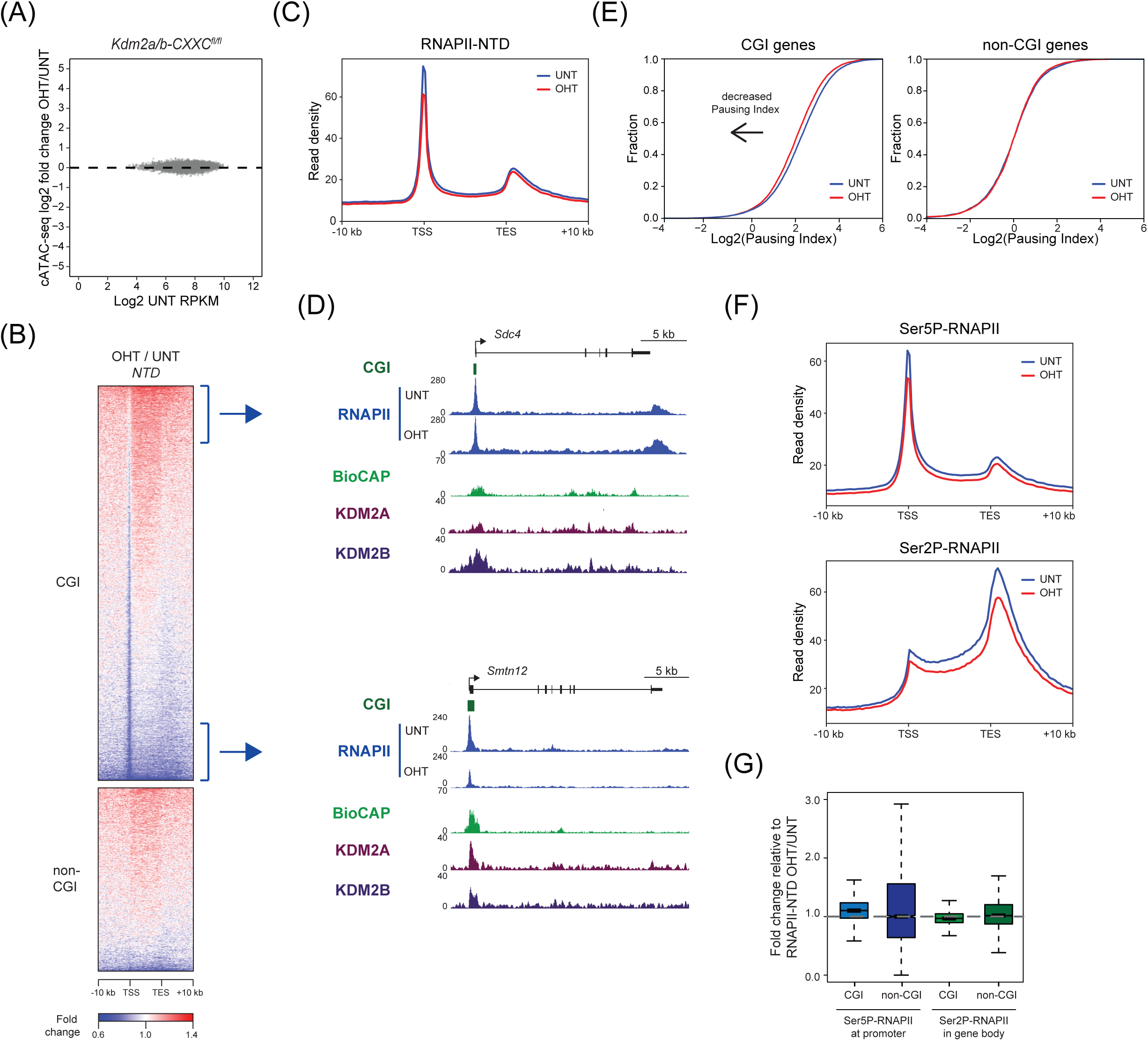
KDM2 proteins regulate RNAPII occupancy but not chromatin accessibility at CGI-associated gene promoters. (A) An MA-plot showing log2 fold change in the accessibility (cATAC-seq) of CGI-associated gene promoters in K*dm2a/b-CXXC*^*fl/fl*^ mESCs following tamoxifen treatment. No promoters significantly changed in accessibility (p-adj < 0.05 and > 1.4-fold). (B) A heatmap of the fold change in RNAPII ChIP-seq signal following tamoxifen treatment of K*dm2a/b-CXXC*^*fl/fl*^ mESCs, for CGI-associated (n=14106) and non-CGI-associated (n=6527) gene promoters. (C) A metaplot showing RNAPII enrichment at CGI-associated genes before (UNT) and after tamoxifen treatment (OHT) *Kdm2a/b-CXXC*^*fl/fl*^ mESCs. (D) Genomic snapshots showing RNAPII occupancy before (UNT) and after tamoxifen treatment (OHT) of *Kdm2a/b-CXXC*^*fl/fl*^ mESCs, for (above) a representative gene that retains RNAPII at the promoter and increases in RNAPII occupancy throughout the gene body, and (below) a representative gene that decreases in RNAPII at both promoter and gene body regions. BioCAP and KDM2A and KDM2B ChIP-seq signal are shown for reference (Farcas et al. 2012; Long et al. 2013b; Blackledge et al. 2014). (E) Empirical cumulative density function (ECDF) plots of RNAPII pausing index for CGI-associated (left) or non-CGI-associated (right) genes, before (UNT) and after tamoxifen treatment (OHT) of *Kdm2a/b-CXXC*^*fl/fl*^ mESCs. (F) Metaplots showing Ser5P-RNAPII (upper panel) or Ser2P-RNAPII (lower panel) enrichment at CGI-associated genes before (UNT) and after tamoxifen treatment (OHT) of *Kdm2a/b-CXXC*^*fl/fl*^ mESCs. (G) A boxplot showing the fold change in Ser5P-RNAPII at gene promoters (left) or Ser2P-RNAPII at gene bodies (right) following tamoxifen treatment of *Kdm2a/b-CXXC*^*fl/fl*^ mESCs, normalised to RNAPII-NTD signal. The fold changes for CGI-associated and non-CGI-associated genes are shown.

To examine this possibility in more detail, we carried out cChIP-seq for RNAPII before and after removal of KDM2 proteins. This revealed on average a widespread decrease in RNAPII occupancy at the TSSs of CGI-associated genes (Figure 6B,C). In the gene body, alterations in RNAPII occupancy appeared to be related to the level of RNAPII reduction at the gene promoter. Genes that retained promoter associated RNAPII showed increased RNAPII in the gene body and promoters showing reduced RNAPII levels also had moderately reduced RNAPII in the gene body (Figure 6D). Importantly, these effects were restricted to CGI-associated genes, in agreement with the function of KDM2A/B at CGIs. To examine in more detail the nature of the defects in RNAPII function at CGI-associated gene promoters, we calculated the RNAPII pausing index, which is often used a proxy for RNAPII pause-release (Supplementary Figure 6B). This showed a modest but clear decrease in pausing index when KDM2 proteins were removed, and importantly this effect was not observed for non-CGI-associated genes (Figure 6E). This suggests that removal of KDM2 proteins may contribute to an increased rate of RNAPII pause release from CGI-associated gene promoters. A comparison of the changes in RNAPII occupancy with the gene expression changes following KDM2 removal revealed a moderate positive correlation (Supplementary Figure 6C), such that the genes that most strongly increased in expression retained RNAPII at their promoter and had elevated RNAPII occupancy throughout the gene body (Supplementary Figure 6D,E). These effects on RNAPII could be explained by an increase in the rate of transcription initiation at genes that show large increases in expression which, when combined with an increased rate of pause release, results in the accumulation of RNAPII throughout the gene body and increases in transcript levels. Importantly, the change in the distribution of Ser5-phosphorylated RNAPII, which is enriched at promoter regions, and Ser2-phosphorylated RNAPII, which is enriched throughout gene bodies, resembled that of total RNAPII (Supplementary Figure 6F). There was no obvious shift in the position of either Ser2-or Ser5-phosphorylated RNAPII enrichment throughout CGI genes (Figure 6F), and only minor changes in the relative enrichment of Ser5-or Ser2-phosphorylated RNAPII compared to total RNAPII at CGI gene promoters and gene bodies, respectively (Figure 6G). Together our findings suggest that KDM2 proteins play a role in regulating RNAPII activity at gene promoters, potentially by limiting initiation and pause release to constrain productive transcription from these regions of the genome.

## DISCUSSION

Chromatin modifying complexes are thought to play central roles in regulating gene expression through their enzymatic activities. Yet, for most of these complexes, the importance of their histone modifying activities in gene regulation remains to be tested. KDM2 histone demethylases have been proposed to contribute to an H3K36me2-depleted and transcriptionally permissive chromatin state at CGI-associated gene promoters. Alternatively, they have also been suggested to contribute to gene repression in some specific instances. However, the extent to which KDM2 proteins regulate gene expression and how this is related to their H3K36me2 demethylase activity has remained untested. Here, using combinatorial genetic perturbation and detailed genome-wide approaches, we discover that the histone demethylase activity of KDM2 proteins contributes modestly to the H3K36me2 depletion at CGI-associated gene promoters (Figure 1) and has minimal effects on gene expression (Figure 2). In contrast, using calibrated gene expression analysis we discover an unexpectedly widespread histone demethylase-independent role for KDM2 proteins in constraining the expression of CGI-associated genes (Figure 3). Importantly, repression by KDM2 proteins is not limited to polycomb target genes, which are known to be regulated by the KDM2B-PRC1 complex (Figure 4). Nevertheless, we find that KDM2B plays the predominant role in repressing expression, while KDM2A appears to contribute at a subset of genes (Figure 5). Finally, the effects of KDM2 proteins on gene expression are not mediated through changes in chromatin accessibility, but instead KDM2 proteins appear to play a role in constraining RNAPII occupancy and possibly pause release at CGI-associated gene promoters to limit transcription (Figure 6). Together, this demonstrates that KDM2 proteins regulate gene expression independently of their histone demethylase activity and through mechanisms that appear to regulate RNAPII function at CGI-associated gene promoters. These discoveries reveal an interesting new chromatin modification-independent role for CGIs and the KDM2 proteins in constraining gene expression.

Our understanding of how histone modification states are specified and regulated remains poorly understood. In the context of histone H3K36, NSD1-3 and ASH1L, the main H3K36me1/2 methyltransferases, can associate with gene promoters and genic regions, and H3K36me2 blankets most of the genome (Gregory et al. 2007; Lucio-Eterovic et al. 2010; Kuo et al. 2011; Rahman et al. 2011; Ram et al. 2011; Shen et al. 2015). However, our genome-wide profiling of H3K36me2 reveals that the bodies of highly transcribed genes and CGI-associated gene promoters are exceptions to this, being uniquely depleted of H3K36me2. This suggests that mechanisms must function to shape H3K36me2 at distinct regions of the genome. The depletion of H3K36me2 in highly transcribed gene bodies is likely due to conversion to H3K36me3 by the SETD2 protein, which interacts with RNAPII and functions as an H3K36 trimethyltransferase in gene bodies during transcriptional elongation (Strahl et al. 2002; Krogan et al. 2003; Li et al. 2003; Schaft et al. 2003; Xiao et al. 2003; Kizer et al. 2005; Li et al. 2005; Sun et al. 2005; Edmunds et al. 2008). We and others had previously proposed that the H3K36me2-depleted state at CGI-associated gene promoters relies on KDM2 proteins to actively remove H3K36me2 from these regions. Now, using a cell system where we can induce the removal of KDM2 demethylase activity, we show that KDM2 enzymes contribute modestly to depletion of H3K36me2 at CGI-associated gene promoters. This suggests that the activity of NSD/ASH1L may be inhibited, or that other histone demethylases may compensate for the loss of KDM2 enzymes, at these regions of the genome. The latter of these two possibilities seems the most likely, as KDM4A-C demethylases catalyse the removal of H3K36me2/3 (Cloos et al. 2006; Fodor et al. 2006; Klose et al. 2006; Whetstine et al. 2006) and also associate broadly with gene promoters (Pedersen et al. 2014; Pedersen et al. 2016). Therefore, determining whether the H3K36me2-depleted state at CGI-associated gene promoters results from active removal of this modification and contributes to gene regulation awaits combinatorial removal of KDM2 and KDM4 demethylase activity.

When studied in the context of individual genes, KDM2A and KDM2B have been proposed to function in both gene activation and repression. However, which of these activities is most prevalent and whether these paralogous proteins function together to achieve appropriate gene regulation have remained unknown. Using combinatorial inducible genetic perturbation strategies and calibrated RNA-seq we now reveal that KDM2 proteins function primarily as repressors of gene expression and elicit their effects via a demethylase-independent mechanism. Gene repression by KDM2 proteins is remarkably widespread but largely restricted to CGI-associated genes, in agreement with the occupancy of KDM2 proteins at these regions of the genome through their ZF-CxxC DNA binding domain. However, lowly expressed genes were more susceptible to increases in gene expression when KDM2 proteins were removed. Therefore, KDM2 proteins may function to generically constrain transcription from CGI-associated gene promoters, but only function to counteract low-level activation signals. In agreement with this suggestion, CGI-associated genes that are already highly expressed are largely unaffected by KDM2 loss, despite the fact that KDM2 proteins occupy their promoters. In the context of these observations, we propose that the repressive activity of KDM2 proteins may effectively create a CGI-imposed barrier to gene activation which protects against low-level or inappropriate gene activation signals. This could be particularly important in the context of cellular differentiation where excessive gene expression noise or precocious gene activation may have deleterious consequences for the highly orchestrated cascade of gene expression events that lead to appropriate acquisition of new cell fates.

Gene repression by the KDM2 proteins occurs independently of their JmjC domain and histone demethylase activity, raising the interesting question of how they repress gene expression. By examining the binding of RNAPII and its modified forms throughout the genome following loss of KDM2 proteins, we discover that there is a widespread reduction in RNAPII occupancy at CGI-associated gene promoters. This suggests that one activity of KDM2 proteins at CGIs may be to constrain productive transcription, perhaps through a process that directly regulates RNAPII pause release. It is intriguing to note that, on average, RNAPII occupancy moderately decreased in gene bodies following loss of KDM2 proteins, despite a widespread increase in gene expression. This raises the possibility that there are further alterations to RNAPII behaviour, such as elongation rate, following KDM2 removal. Based on these observations, an important area of future work will be to examine the mechanisms by which KDM2 proteins affect RNAPII activity and to determine how direct this is.

We speculate that one mechanism by which KDM2 proteins could potentially modulate RNAPII-dependent transcription processes is through ubiquitination. This is because, in addition to their JmjC domains, KDM2 proteins also encode FBOX and LRR domains. The FBOX binds a protein called SKP1, and we and others have previously shown that KDM2A and KDM2B both interact with SKP1 (Gearhart et al. 2006; Koyama-Nasu et al. 2007; Farcas et al. 2012; Tan et al. 2013). SKP1 is a central component of SCF-type E3 ubiquitin ligase complexes (Cardozo and Pagano 2004), while FBOX-containing proteins are thought to confer substrate specificity for SCF complexes through additional domains such as the LRR domain (Ho et al. 2006). This suggests that KDM2 proteins might identify target proteins for ubiquitylation. Indeed, KDM2B was reported to ubiquitylate the transcription factor c-Fos, leading to its degradation by the proteasome (Han et al. 2016). KDM2A has also been proposed to possess E3 ligase activity, as its overexpression stimulates 53BP1 ubiquitylation (Bueno et al. 2018). The specificity of these putative KDM2 E3 ubiquitin ligase complexes remains to be investigated. However, given that KDM2 proteins act broadly to repress gene expression and may regulate RNAPII activity, one might envisage that KDM2 proteins could regulate a component of the core transcriptional machinery or another general modulator of gene transcription. Therefore, in future work it will be interesting to explore whether KDM2 proteins have a role in proteostasis at CGIs and to understand whether this contributes to their function in the repression of gene expression and the regulation of RNAPII activity.

In conclusion, we discover that KDM2 proteins are CGI-specific transcriptional repressors that appear to function to constrain low-level gene activation signals. Interestingly, DNA situated in CGIs is known to be highly accessible, differentiating it from much of the rest of the genome. It has been proposed that this accessibility highlights the location of gene regulatory elements within large and complex vertebrate genomes, and allows transcriptional regulators and the transcriptional machinery to more easily access the underlying DNA and enable gene expression. However, an unintended consequence of this CGI-associated accessibility may be that it renders these regions susceptible to low-level and potentially inappropriate gene activation signals. We speculate that, in response to this potentially deleterious side effect of CGI accessibility, KDM2 proteins may have evolved to bind CGIs and constrain transcription. Indeed, we show that loss of KDM2 proteins does not affect accessibly at CGIs but does broadly affect gene expression and RNAPII occupancy. Therefore we propose that CGIs create an appropriate balance of transcriptionally permissive and restrictive activities to help control gene expression.

## MATERIALS AND METHODS

### Cell culture

Mouse embryonic stem cells (mESCs) were cultured on gelatine-coated dishes at 37°C and 5% CO_2_, in DMEM (Life Technologies) supplemented with 15% fetal bovine serum (Labtech), 2mM L-glutamine (Life Technologies), 0.5 mM beta-mercaptoethanol (Life Technologies), 1x non-essential amino acids (Life Technologies), 1x penicillin-streptomycin (Life Technologies), and 10 ng/ml leukemia-inhibitory factor. Conditional mESC lines were treated with 800 nM 4-hydroxytamoxifen (Sigma) for 96 hours to induce KDM2-LFs knockout (*Kdm2a/b-JmjC*^*fl/fl*^) or ZF-CxxC domain deletion (*Kdm2a/b-CXXC*^*fl/fl*^, *Kdm2a-CXXC*^*fl/fl*^, *Kdm2b-CXXC*^*fl/fl*^).

Human HEK293T cells used for RNAPII cChIP-seq were grown at 37°C and 5% CO_2_ in DMEM supplemented with 10% fetal bovine serum, 2 mM L-glutamine, 0.5 mM beta-mercaptoethanol and 1x penicillin-streptomycin. *D. melanogaster* S2 (SG4) cells used for cnRNA-seq were grown adhesively at 25°C in Schneider’s Drosophila Medium (Life Technologies), supplemented with 1x penicillin-streptomycin and 10% heat-inactivated fetal bovine serum.

### Generation of the *Kdm2a/b-JmjC*^*fl/fl*^ mESC line

To generate the *Kdm2a/b-JmjC*^*fl/fl*^ mESC line, a loxP site was inserted upstream of the critical JmjC domain-encoding exon(s) in the *Kdm2a* and *Kdm2b* genes (exon 8 for *Kdm2a*, and exons 7-8 for *Kdm2b*), and FRT flanked PGK-neo and a second loxP site was inserted downstream of the critical exon(s). Targeting vectors were generated from bacterial artificial chromosomes containing the target mouse genomic regions using the Double Red recombination method, as previously described (Suzuki and Nakayama 2011). Linearized targeting vectors were introduced into M1 mESCs by electroporation (GenePulser, Bio-Rad). mESC colonies were isolated and expanded, and the genomic DNA of each clone was purified. Homozygous loxP targeting was verified by sequencing of the genomic region surrounding the loxP sites. Targeted ES cells were injected into mouse blastocysts to generate chimeric mice. The *Kdm2a/b-JmjC*^*fl/fl*^ line was generated by removal of the PGK-neo marker gene by mating the targeted mice with mice expressing FLP recombinase. These *Kdm2a/b-JmjC*^*fl/fl*^ mice were further mated with mice harboring the ROSA26-CreErt2 locus to generate *Kdm2a/b-JmjC*^*fl/fl*^*:ROSA26-CreErt2*^*+/-*^ mice, from which the *Kdm2a/b-JmjC*^*fl/fl*^ mESCs used in this study were derived.

### Generation of *Kdm2a-CXXC*^*fl/fl*^ and *Kdm2a/b-CXXC*^*fl/fl*^ mESC lines

Conditional *Kdm2a-CXXC*^*fl/fl*^ and *Kdm2a/b-CXXC*^*fl/fl*^ mESC lines were generated by using CRISPR-mediated genome editing to insert parallel loxP sites flanking exon 14 of the *Kdm2a* gene in *Rosa26::CreERT2* or *Kdm2b-CXXC*^*fl/fl*^ mESCs, respectively (Blackledge et al. 2014). Targeting constructs encoding the loxP sequence flanked by 150 bp homology arms and carrying a mutated PAM sequence to prevent retargeting by the Cas9 enzyme were purchased from GeneArt (ThermoFisher). The pSpCas9(BB)-2A-Puro(PX459)-V2.0 vector was obtained from Addgene (#62988). sgRNAs were designed using the CRISPOR online tool (http://crispor.tefor.net/crispor.py) and were cloned into the vector as previously described (Ran et al. 2013). First, the upstream loxP site was targeted. *Rosa26::CreERT2* or *Kdm2b-CXXC*^*fl/fl*^ mESCs were transiently co-transfected with 1 μg of Cas9-sgRNA plasmid and 3.5 μg of targeting construct using Lipofectamine 3000 (ThermoFisher). The day after transfection, cells were passaged at a range of densities and subjected to puromycin selection (1 μg/ml) for 48 hours. Individual clones were isolated and PCR-screened. A correctly targeted homozygous clone was then used to target the downstream loxP site using the same transfection protocol and screening strategy. Correct loxP targeting was verified by sequencing of the genomic region surround the loxP sites, and clones were analysed by both RT-qPCR and western blot to confirm loss of the ZF-CxxC domain in response to tamoxifen treatment.

### Protein extracts and immunoblotting

For nuclear extraction, mESCs were washed with PBS then resuspended in 10 volumes of Buffer A (10mM Hepes pH 7.9, 1.5mM MgCl_2_, 10mM KCl, 0.5mM DTT, 0.5mM PMSF, and 1x PIC (Roche)) and incubated on ice for 10 min. Cells were recovered by centrifugation at 1500 g for 5 min, resuspended in 3 volumes of Buffer A supplemented with 0.1% NP-40 and incubated on ice for 10 min. The released nuclei were recovered by centrifugation at 1500 g for 5 min and resuspended in 1 pellet volume of Buffer B (5mM Hepes pH 7.9, 26% glycerol, 400mM NaCl, 1.5mM MgCl2, 0.2mM EDTA, 0.5mM DTT and 1x PIC). After 1 hour of rotation at 4°C, the suspension was pelleted at 16,000 g for 20 min and the supernatant taken as nuclear extract.

For histone extraction, mESCs were washed with RSB (10mM Tris HCl pH 7.4, 10mM NaCl, 3mM MgCl_2_ and 20mM NEM), then resuspended in RSB buffer supplemented with 0.5% NP-40 and incubated on ice for 10 min to allow cell lysis. Following centrifugation at 500 g for 5 min, the nuclear pellet was incubated in 2.5mM MgCl_2_, 0.4M HCl and 20mM NEM on ice for 20 min. After centrifugation at 16,000 g for 20 min, histones were precipitated from the supernatant on ice with 25% TCA for 30 min. Histones were recovered by centrifugation at 16,000 g for 15 min, and the pellet was washed twice in acetone. The histone pellet was resuspended in 1x SDS loading buffer and boiled at 95°C for 5 min. Any insoluble precipitate was pelleted by centrifugation at 16,000 g for 15 min and the soluble fraction retained as histone extract. Histone concentrations were compared by Coomassie Blue staining following SDS-PAGE. Semi-quantitative western blot analysis of histone extracts was performed using LiCOR IRDye® secondary antibodies and the LiCOR Odyssey Fc system. To measure changes in H3K36 methylation, the signal relative to H4 histone was determined.

### Antibodies

The following antibodies were used in this study: anti-KDM2A (Blackledge et al. 2010), anti-KDM2B (Farcas et al. 2012), anti-BRG1 (EPNCIR111A, Abcam), anti-H3, anti-H3K36me1, anti-H3K36me2 (Blackledge et al. 2010), anti-H3K36me3, anti-H4 (L64C1, Cell Signalling), anti-Rbp1-NTD (D8L4Y, Cell Signalling), anti-Rbp1-CTD-Ser5P (D9N5I, Cell Signalling), anti-Rbp1-CTD-Ser2P (E1Z3G, Cell Signalling). Anti-H3 and anti-H3K36me antibodies were prepared in-house by rabbit immunisation with synthetic peptides (PTU/BS Scottish National Blood Transfusion Service), and antibodies were purified on peptide affinity columns.

### Preparation of chromatin

For KDM2A/B ChIP, 5×10^7^ mESCs were resuspended in PBS and crosslinked in 2 mM disuccinimidyl glutarate (Thermo Scientific) for 45 min at 25°C with gentle rotation, then in 1% formaldehyde for 12.5 min (methanol-free, Life Technologies). Reactions were quenched by addition of 125 mM glycine, and crosslinked cells were resuspended in lysis buffer (50mM HEPES-KOH pH 7.9, 140mM NaCl, 1mM EDTA, 10% glycerol, 0.5% NP40, 0.25% TritonX-100 and 1x PIC) and rotated for 10 min at 4°C. The released nuclei were washed (10 mM Tris-HCl pH 8.0, 200mM NaCl, 1mM EDTA, 0.5mM EGTA and 1x PIC) for 5 min at 4°C, and the nuclear pellet resuspended in 1 ml sonication buffer (10mM Tris HCl pH 8.0, 100mM NaCl, 1mM EDTA, 0.5mM EGTA, 0.1% sodium deoxycholate, 0.5% N-lauroylsarcosine and 1x PIC).

For histone ChIP, 1×10^7^ mESCs were crosslinked for 10 min in 1% formaldehyde. Reactions were quenched by addition of 125 mM glycine. The released nuclei washed twice in PBS, then resuspended in lysis buffer (1% SDS, 10mM EDTA, 50mM Tris HCl pH 8.1 and 1x PIC) and incubated on ice for 30 min.

For RNAPII ChIP, 5×10^7^ mESCs were resuspended in PBS and mixed with 4×10^6^ HEK293T cells. Cells were crosslinked for 10 min in 1% formaldehyde. Reactions were quenched by addition of 150 mM glycine, and the crosslinked cells resuspended in FA-lysis buffer for 10 min (50mM HEPES pH 7.9, 150mM NaCl, 2mM EDTA, 0.5mM EGTA, 0.5% NP40, 0.1% sodium deoxycholate, 0.1% SDS, 10mM NaF, 1mM AEBSF, 1x PIC).

Chromatin was sonicated using a BioRuptor Pico sonicator (Diagenode), shearing genomic DNA to approximately 0.5 kb. Following sonication, TritonX-100 was added to chromatin used for KDM2A/B ChIP to a final concentration of 1%.

### Chromatin immunoprecipitation and sequencing

Sonicated chromatin was diluted 10-fold in ChIP dilution buffer (1% Triton-×100, 1 mM EDTA, 20mM TrisHCl pH 8, 150mM NaCl and 1x PIC) for KDM2A/B or histone ChIP, or in FA-lysis buffer for RNAPII ChIP. Chromatin was pre-cleared for 1 hour with either protein A magnetic Dynabeads (Invitrogen, for KDM2A/B ChIP) or protein A agarose beads (Repligen, for histone or RNAPII ChIP) blocked with 1 mg/ml BSA and 1 mg/ml yeast tRNA. For each ChIP reaction, 150 μg chromatin (KDM2A/B), 300 μg chromatin (RNAPII) or chromatin corresponding to 1×10^5^ cells (histone ChIP) was incubated overnight with the appropriate antibody: anti-KDM2A (2.4 μl), anti-KDM2B (2 μl), anti-H3 (15 μl) anti-H3K36me2 (15 μl), anti-Rbp1-NTD (15 μl), anti-Rbp1-CTD-Ser5P (12.5 μl), anti-Rbp1-CTD-Ser2P (12.5 μl).

Antibody-bound chromatin was isolated using blocked protein A agarose (histone/ RNAPII ChIP) or magnetic beads (KDM2A/B ChIP) for 2 hours at 4°C. For histone or KDM2A/B ChIP, washes were performed with low salt buffer (0.1% SDS, 1% TritonX-100, 2 mM EDTA, 20 mM Tris-HCl pH 8, 150 mM NaCl), high salt buffer (0.1% SDS, 1% TritonX-100, 2 mM EDTA, 20 mM Tris-HCl pH 8, 500 mM NaCl), LiCl buffer (250mM LiCl, 1% NP40, 1% sodium deoxycholate, 1 mM EDTA, 10 mM Tris-HCl pH 8) and two washes with TE buffer (10 mM Tris-HCl pH 8, 1 mM EDTA). For RNAPII ChIP, washes were performed with FA-Lysis buffer, FA-Lysis buffer containing 500mM NaCl, DOC buffer (250mM LiCl, 0.5% NP40, 0.5% sodium deoxycholate, 2 mM EDTA, 10mM Tris-HCl pH 8) and two washes with TE buffer. ChIP DNA was eluted in elution buffer (1% SDS, 100mM NaHCO_3_) and crosslinks reversed overnight at 65°C with 200 mM NaCl and 2 μl RNase A (Sigma). A matched input sample (corresponding to 10% of original ChIP reaction) was treated identically. The following day, samples were treated with 20 μg/ml Proteinase K (Sigma) for 2 hours at 45°C and purified using the ChIP DNA Clean and Concentrator Kit (Zymo Research).

ChIP-seq libraries for both ChIP and input samples were prepared using the NEBNext Ultra DNA Library Prep Kit for Illumina (NEB), following the manufacturer’s guidelines and using NEBNext Multiplex Oligos. The average size and concentration of libraries were determined using the 2100 Bioanalyzer High Sensitivity DNA Kit (Agilent) and qPCR with SensiMix SYBR (Bioline) and KAPA Illumina DNA standards (Roche). Libraries were sequenced using the Illumina NextSeq 500 platform in biological triplicate or quadruplicate with 40 bp paired-end reads.

### Calibrated nuclear RNA-sequencing and ATAC-sequencing (cnRNA-seq and cATAC-seq)

To isolate nuclei for cnRNA-seq and cATAC-seq, 10^7^ mESCs were mixed with 2.5×10^6^ Drosophila SG4 cells in PBS. Cells were lysed in 1 ml HS lysis buffer (0.05% NP40, 50mM KCl, 10 mM MgSO4.7H20, 5mM HEPES, 1mM PMSF, 3mM DTT, 1x PIC). Nuclei were recovered by centrifugation at 1000 g for 5min and washed three times in 1 ml resuspension buffer (10mM NaCl, 10mM Tris pH 7.4, 3mM MgCl_2_). Nuclear integrity was assessed using 0.4% trypan blue staining (ThermoFisher).

Nuclear RNA was prepared from 4×10^6^ nuclei using TRIzol reagent according to the manufacturer’s protocol (Invitrogen), then treated with the TURBO DNA-free kit (ThermoFisher). nRNA quality was assessed using the 2100 Bioanalyzer RNA 6000 Pico kit (Agilent), then nRNA was depleted of rRNA using the NEBNext rRNA depletion kit and the depletion efficiency evaluated using the Bioanlayzer RNA 6000 Pico kit. RNA-seq libraries were prepared using the NEBNext Ultra Directional RNA-seq kit, and library size and concentration was determined as described for ChIP libraries. Libraries were sequenced using the Illumina NextSeq 500 platform in biological triplicate or quadruplicate using 80 bp paired-end reads.

Chromatin accessibility was assayed using an adaptation of the assay for transposase accessible-chromatin (ATAC)-seq (Buenrostro et al. 2013) as previously described (King and Klose 2017), using 5×10^5^nuclei from the same preparation used for the purification of nuclear RNA. Genomic DNA was also purified from an aliquot of the same preparation of nuclei by phenol-chloroform extraction and tagmented with Tn5, to control for sequence bias of the Tn5 transposase and to determine the exact mouse/fly mixing ratio for each individual sample.

ATAC-seq and input gDNA libraries were prepared by PCR amplification using custom-made Illumina barcodes (Buenrostro et al. 2013) and the NEBNext High-Fidelity 2X PCR Master Mix. Libraries were purified with two rounds of Agencourt AMPure XP bead cleanup (Agencourt, 1.5X bead:sample ratio). Library size and concentration were determined as described for ChIP libraries. Libraries were sequenced using the Illumina NextSeq 500 platform in biological triplicate or quadruplicate using 80 bp paired-end reads.

### Digital droplet PCR

For digital droplet PCR (ddPCR), total RNA was prepared from 10^6^ mESCs using the RNeasy mini plus kit including gDNA eliminator columns (QIAGEN). RPT low retention tips (Starlab) were used throughout the ddPCR protocol to increase pipetting accuracy. Purified RNA was eluted in 30 μl elution buffer, 7 μl was diluted with 8 μl water, and 4 μl of this dilution was reverse transcribed using the imProm-II system with random hexamer primers and RNasin ribonuclease inhibitor (Promega). The generated cDNA was diluted with 300 ul nuclease-free water. ddPCR primers were designed using Primer 3 Plus (Untergasser et al. 2012) with BioRad recommended settings: 3.8mM divalent cations, 0.8mM dNTPs, 80-120 bp product, 60-61°C melting temperature. Their efficiency was tested using a serial dilution curve of cDNA by standard SYBR qPCR. ddPCR reactions were prepared in 96-well PCR plates, and contained 12.5 μl 2x QX200 ddPCR EvaGreen Supermix (BioRad), 0.32 μl each of forward and reverse primers (10 μM), 7 μl diluted cDNA and 4.86 μl nuclease-free water. This 25 μl reaction was mixed by pipetting, then 22 μl was transferred to a semi-skirted 96-well PCR plate (Eppendorf) and used for droplet generation with an AutoDG droplet generator (BioRad). Droplets were collected in a semi-skirted PCR plate, which was then sealed using a PX1 PCR plate sealer (BioRad). PCR was performed using a C1000 Touch thermal cycler (BioRad) with a 2°C/s ramp rate: 5 min at 95°C followed by 40 cycles of denaturation at 95°C for 30 s and annealing/extension at 60°C for 60 s, then signal stabilisation at 4°C for 5 min followed by 90°C for 5 min. Droplets were sorted into PCR-positive and PCR-negative fractions according to their fluorescence using the QX200 droplet reader (BioRad). QuantaLife software (BioRad) was used to calculate the absolute concentration of template cDNA in the ddPCR reaction.

### Data processing and normalisation of massively-parallel sequencing

For cATAC-seq and RNAPII cChIP-seq (including sequencing of input gDNA), reads were aligned to concatenated mouse and spike-in genomes (mm10+dm6 or mm10+hg19) using Bowtie 2 with the ‘-- no-mixed’ and ‘-- no-discordant’ options (Langmead and Salzberg 2012). For histone ChIP-seq, reads were aligned to the mouse mm10 genome as above. Reads that were mapped more than once were discarded, and PCR duplicates were removed using SAMTools (Li et al. 2009b). For cATAC-seq, reads that mapped to a custom ‘blacklist’ of genomic regions with artificially high counts, including mitochondrial DNA sequences, were also discarded.

For cnRNA-seq, reads were first aligned using Bowtie 2 (with ‘--very-fast’, ‘--no-mixed’ and ‘-- no-discordant’ options) to the concatenated mm10 and dm6 rRNA genomic sequence (GenBank: BK000964.3 and M21017.1) to filter out reads mapping to rRNA. All unmapped reads were aligned to the concatenated mm10+dm6 genome using STAR (Dobin et al. 2013). To improve mapping of intronic sequences, reads that failed to map using STAR were aligned using Bowtie 2, with ‘--sensitive-local’, ‘--no-mixed’ and ‘--no-discordant’ options. PCR duplicates were removed using SAMtools (Li et al. 2009b).

To internally calibrate cnRNA-seq, cATAC-seq and cChIP-seq experiments we spiked a fixed number of control cells into each sample (Drosophila SG4 cells for cnRNA-seq and cATAC-seq, human HEK293T cells for RNAPII cChIP-seq). This spike-in genome was then used to quantitatively compare the gene expression, chromatin accessibility or RNAPII profiles between experimental conditions. For visualisation of cATAC-seq and cChIP-seq data, mm10 reads were randomly subsampled by a factor that reflects the total number of spike-in reads in the same sample, as previously described (Bonhoure et al. 2014; Orlando et al. 2014; Hu et al. 2015). To account for any variation in the exact spike-in cell: mESC mixing ratio between biological replicates, the subsampling factors were additionally corrected according to the ratio of dm6 (or hg19)/mm10 total read counts in the matched input sample. For visualisation of cnRNA-seq data, mm10 reads were randomly subsampled by *Drosophila* normalised size factors calculated using DESeq2 (see below). Histone ChIP-seq libraries were randomly downsampled to achieve the same total number of reads for each individual replicate using SAMtools (Li et al. 2009b).

### Read count quantitation and analysis

To compare replicates, read coverage across regions of interest (gene bodies for cnRNA-seq and ChIP-seq, gene promoters for cATAC-seq) was analysed using deepTools multiBamSummary and plotCorrelation functions (Ramírez et al. 2016). For each condition, biological replicates correlated well with each other (Pearson correlation coefficient > 0.95) and were merged for downstream applications.

Genome coverage tracks were generated using the pileup function from MACS2 (Zhang et al. 2008) for ChIP-seq and ATAC-seq and genomeCoverageBed from BEDtools (Quinlan 2014) for cnRNA-seq and visualised using the UCSC genome browser (Kent et al. 2002). Differential bigwig tracks of H3K36me2 normalised to H3, normalised H3K36me2 signal in tamoxifen-treated versus-untreated cells, or RNAPII signal in tamoxifen-treated versus –untreated cells, were generated from merged bigwig files using the deepTools bigwigCompare function with ‘--operation ratio’ setting (Ramírez et al. 2016). Metaplot and heatmap analyses of read density were performed using the computeMatrix, plotProfile and plotHeatmap deepTools functions (v3.0.1). For ChIP-seq, intervals of interest were annotated with normalised read counts from merged replicates with a custom Perl script using SAMtools, or from differential bigwig files using deepTools computeMatrix with the ‘--outFileNameMatrix’ option. Correlation analyses were performed in R using Spearman correlation and visualised with scatterplots coloured by density using ‘stat_density2d’.

### Differential ATAC-seq and gene expression analyses

DESeq2 (Love et al. 2014) was used with a custom R script to identify significant changes in chromatin accessibility or gene expression. In order to calibrate to the spike-in genome, *Drosophila* reads were first pre-normalised according to the exact dm6/mm10 spike-in ratio derived from the matched input gDNA sample. *Drosophila* read counts were then generated for a set of unique dm6 refGene genes and used to calculate DESeq2 size factors. These size factors were supplied for DESeq2 normalisation of raw mm10 read counts for a custom non-redundant mm10 gene set of 20633 genes. P-adj < 0.05 and fold change > 1.4 thresholds were used to determine significant changes. Log2 fold changes were visualised using MA-plots generated with ggplot2.

### Gene annotation

Non-redundant mouse genes (n = 20633) were classified into non-CGI, PRC and non-PRC categories based on the presence of a non-methylated CGI and RING1B and SUZ12 binding at their promoters. Gene Ontology analysis was performed using DAVID (Huang da et al. 2009). The BP FAT setting and a FDR < 0.1 cut-off were used, and the complete non-redundant mm10 gene set was used as a background.

### Accession numbers

The following previously published datasets were used for analysis: H3K36me3 ChIP-seq (GSE34520) (Brookes et al. 2012), KDM2A ChIP-seq (GSE41267) (Farcas et al. 2012), KDM2B ChIP-seq (GSE55698) (Blackledge et al. 2014), BioCAP (GSE43512) (Long et al. 2013b).

## ACKNOWLEDGEMENTS

We thank Nadezda Fursova for assistance with computational analysis, Emilia Dimitrova, Angelika Feldmann and Neil Blackledge for helpful discussions, and Amy Hughes and Neil Blackledge for critical reading of the manuscript. We are grateful to Amanda Williams at the Department of Zoology in Oxford for sequencing support on the NextSeq 500. Work in the Klose laboratory is supported by the Wellcome Trust, the Lister Institute of Preventive Medicine and the European Research Council. Takashi Kondo and Haruhiko Koseki are supported by the AMED-CREST programme from the Japan Agency for Medical Research and Development.

---

**Figure S1 – Related to Figure 1.**
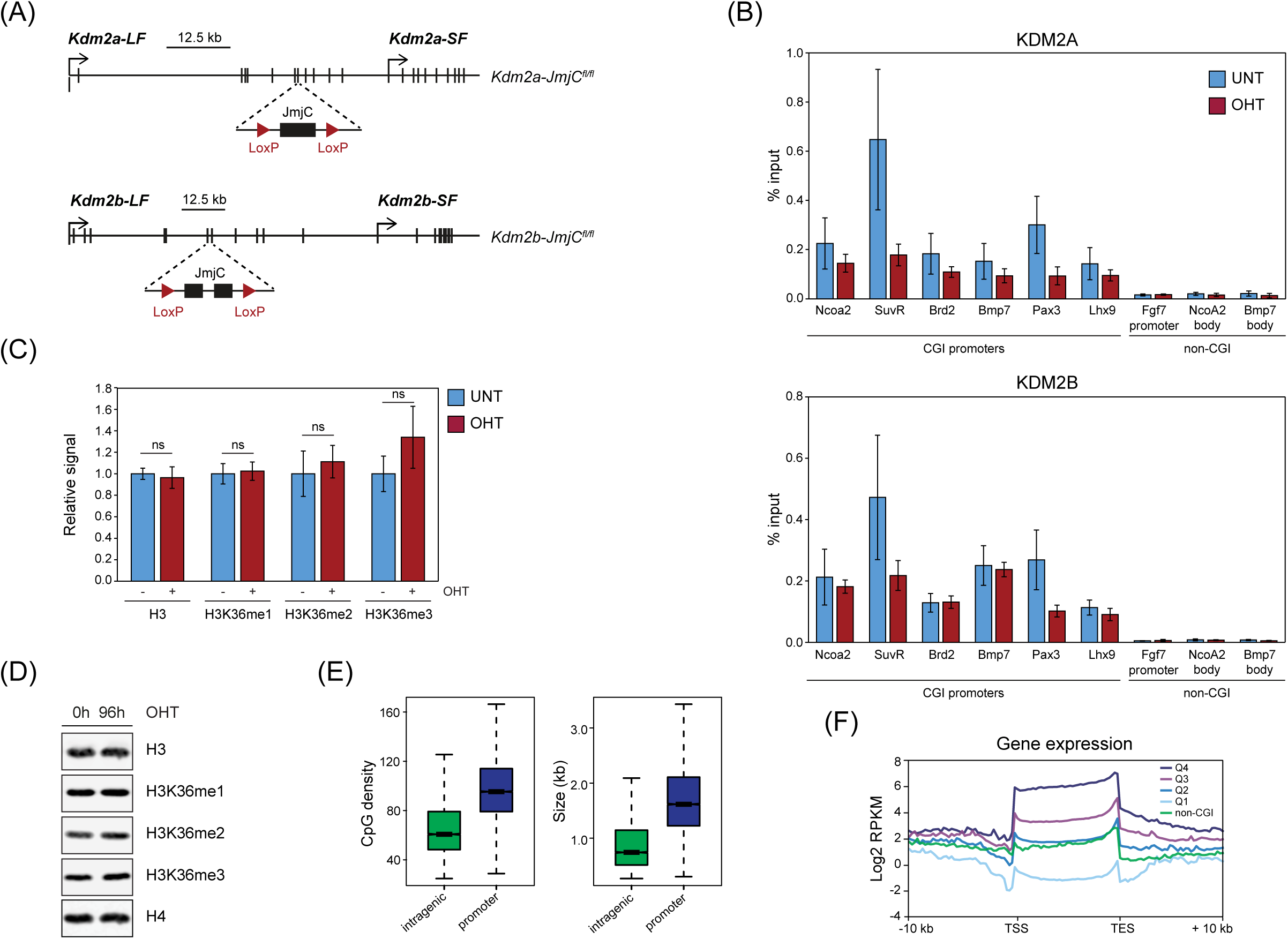
(A) A schematic representation of the *Kdm2a/b-JmjC*^*fl/fl*^ mESC line, in which loxP sites were inserted into the *Kdm2a* and *Kdm2b* genes flanking exons that encode the JmjC domain. (B) ChIP-qPCR analysis showing KDM2A (upper panel) and KDM2B (lower panel) enrichment relative to input in *Kdm2a/b-JmjC*^*fl/fl*^ mESCs before (UNT) and after tamoxifen treatment (OHT). Error bars show standard error of the mean of three biological replicates. (C) Quantitation of western blots of histone extract from *Kdm2a/b-JmjCfl/fl* mESCs, before (UNT) and after 96 hours of tamoxifen treatment (OHT). Signal is normalised to histone H4 and is represented relative to average UNT signal. Error bars show standard deviation of three biological replicates. Significance was tested using a Student’s T-test (non-significant (ns) if p > 0.05). (D) Representative western blots. Histone H4 is shown as a loading control. (E) Boxplots showing CpG density (left) and size (right) of intragenic and promoter-associated CGIs. (F) A metaplot of cnRNA-seq signal in K*dm2a/b-JmjC*^*fl/fl*^ mESCs, for CGI-associated genes separated into quartiles according to their expression level and for non-CGI-associated genes. Genes were scaled to the same length and aligned at their TSS and TES.

**Figure S3 – Related to Figure 3.**
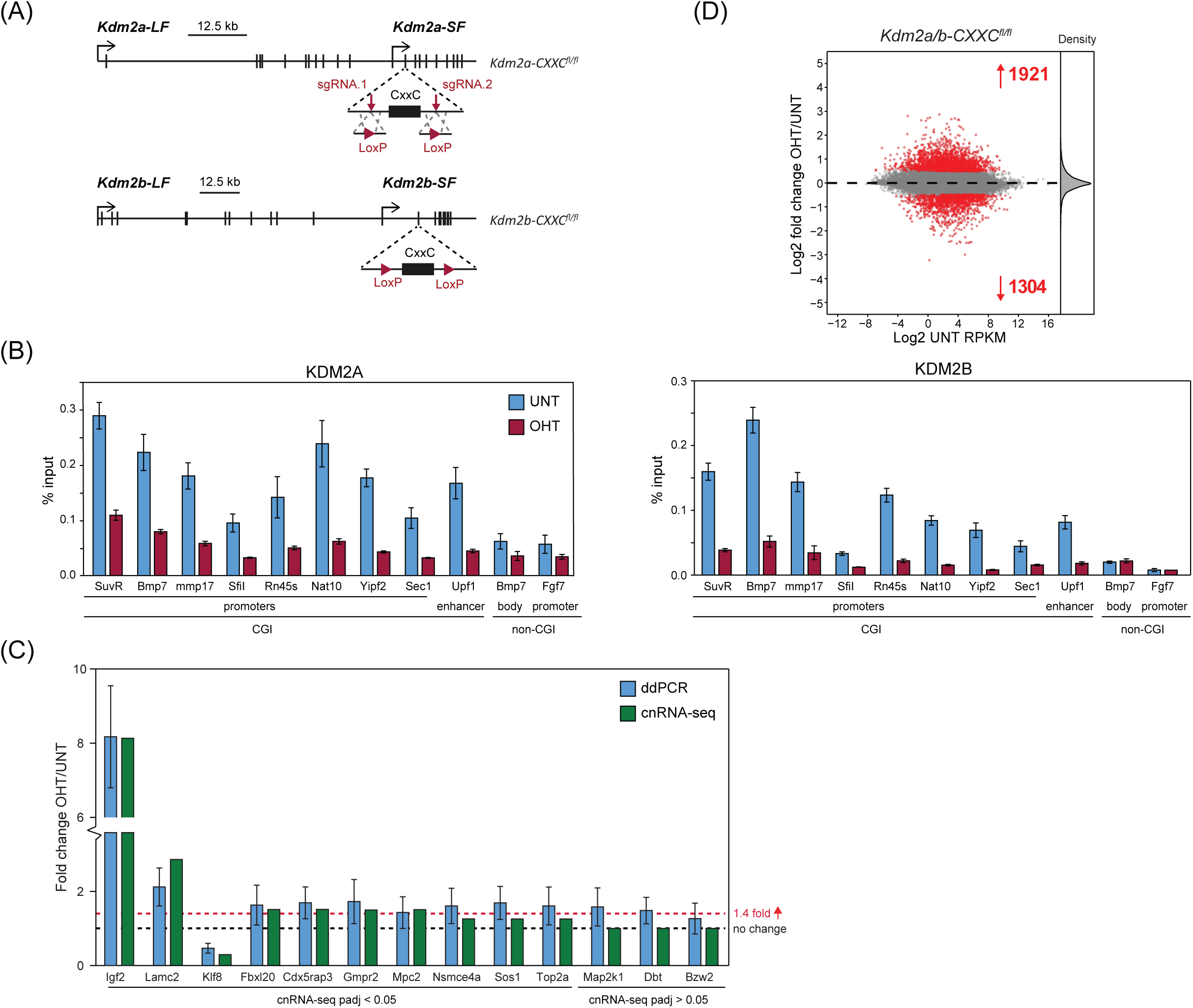
(A) A schematic representation of the *Kdm2a/b-CxxC*^*fl/fl*^ mESC line, in which loxP sites were inserted into the *Kdm2a* and *Kdm2b* genes flanking exons that encode the ZF-CxxC domain. sgRNA.1 and sgRNA.2 indicate the position of CRISPR-mediated loxP insertion. (B) ChIP-qPCR analysis showing KDM2A (left panel) and KDM2B (right panel) enrichment relative to input in *Kdm2a/b-CXXC*^*fl/fl*^ mESCs before (UNT) and after tamoxifen treatment (OHT). Error bars show standard error of the mean of four biological replicates. (C) Digital droplet PCR analysis, showing the fold change in template cDNA concentration following tamoxifen treatment (OHT) of *Kdm2a/b-CXXC*^*fl/fl*^ mESCs (blue). Error bars show standard deviation of three biological replicates. The fold change calculated by cnRNA-seq is shown for comparison (green), and the significance of this change is annotated below the graph. The dashed lines represent no change in expression (black) and the 1.4 fold change threshold used for cnRNA-seq analysis (red). (D) An MA-plot showing log2 fold change in gene expression in *Kdm2a/b-CXXC*^*fl/fl*^ mESCs following tamoxifen treatment (OHT), normalising nRNA-seq data to total library size. The number of genes with significantly increased or decreased expression (p-adj <0.05 and > 1.4-fold) is shown in red and density of gene expression changes is shown on the right.

**Figure S4 – Related to Figure 4.**
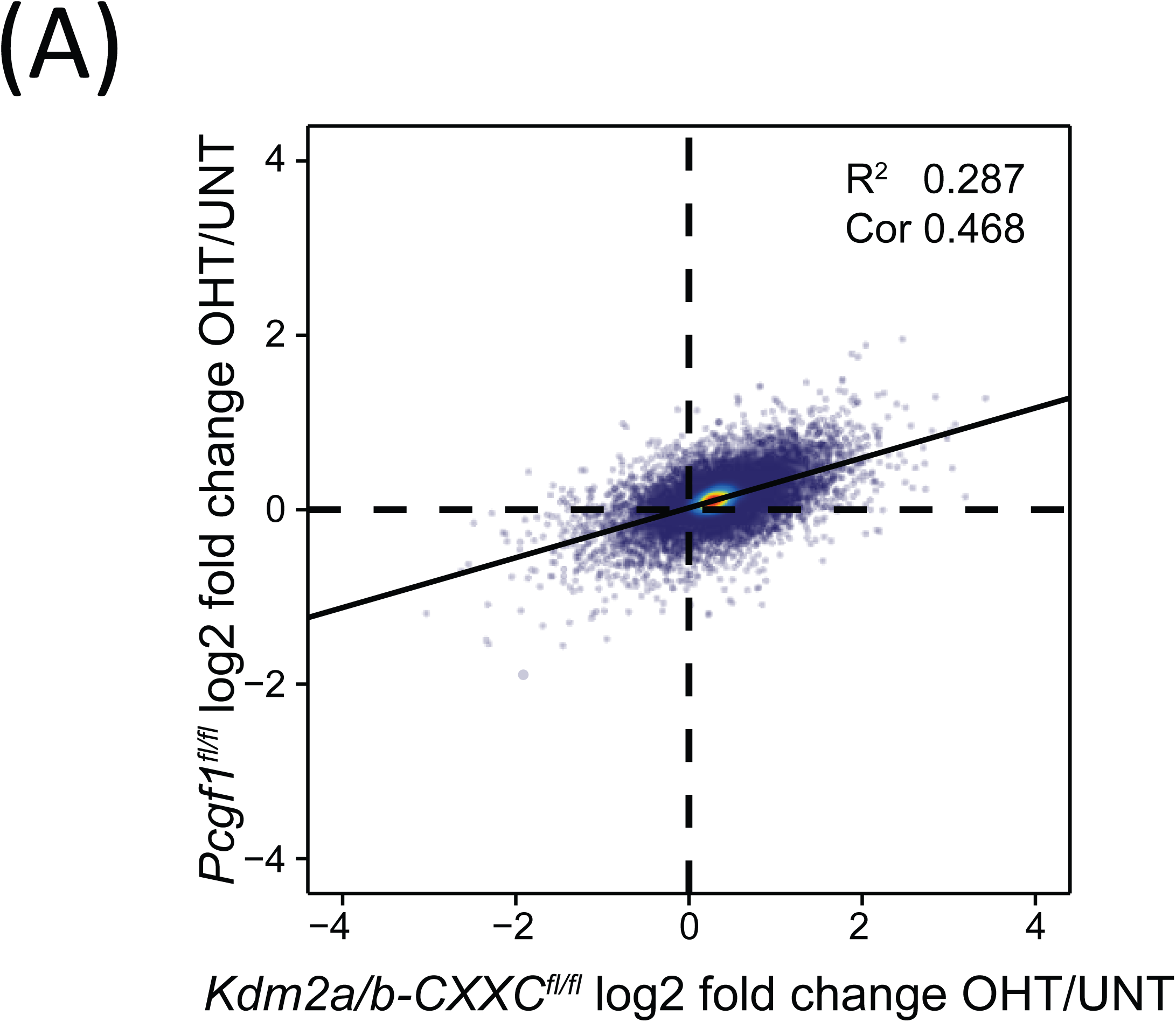
(A) A scatter plot comparing the log2 fold change in gene expression (cnRNA-seq) following tamoxifen treatment of *Kdm2a/b-CXXC*^*fl/fl*^ and *Pcgf1*^*fl/fl*^ mESCs. The solid line shows the linear regression, and the coefficient of determination (R^2^) and Spearman correlation coefficient (Cor) are annotated.

**Figure S5 – Related to Figure 5.**
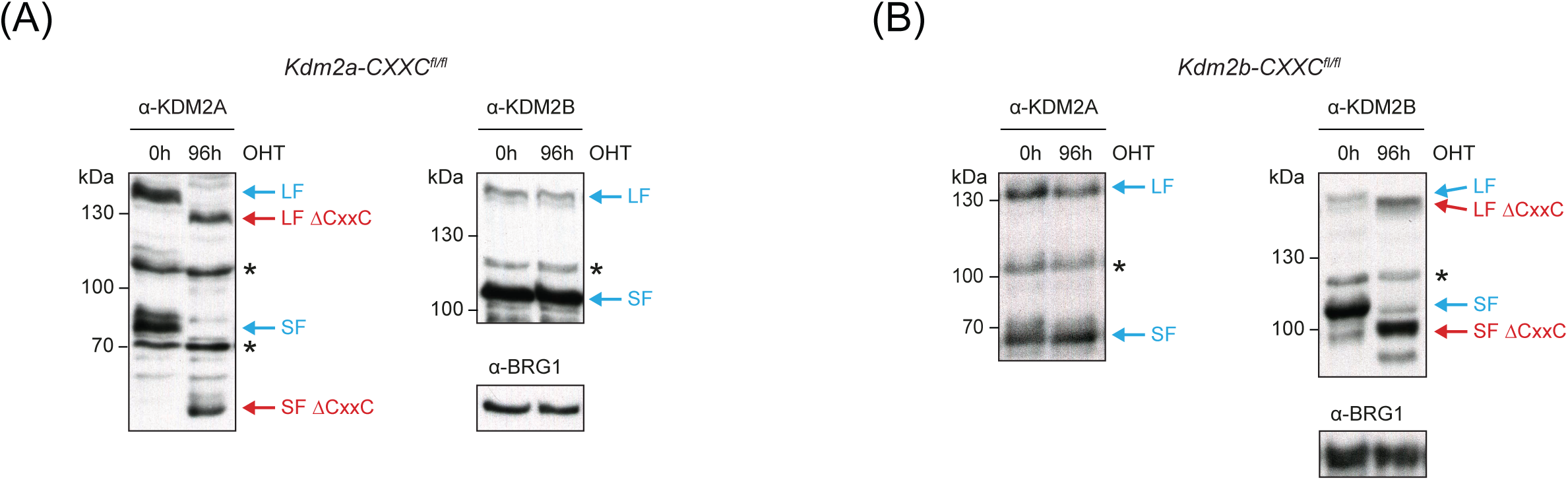
(A) Western blot analysis for KDM2A and KDM2B in K*dm2a-CXXC*^*fl/fl*^ mESCs before (UNT) and after 96 hours of tamoxifen treatment (OHT). BRG1 is shown as a loading control for both blots. Asterisks indicate non-specific bands. (B) As (A) but for K*dm2b-CXXC*^*fl/fl*^ mESCs.

**Figure S6 – Related to Figure 6.**
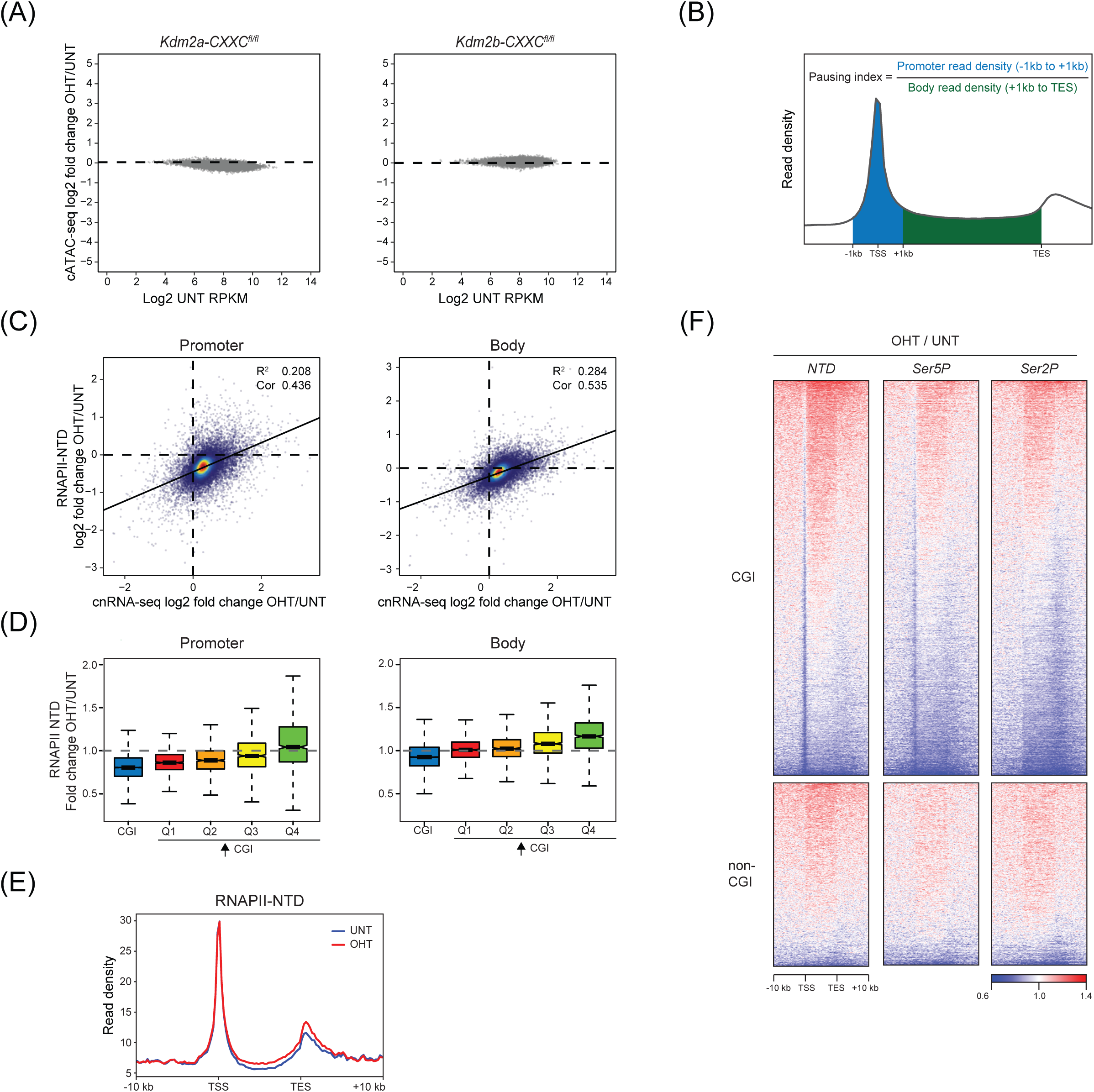
(A) MA-plots showing log2 fold change in the accessibility (cATAC-seq) of CGI-associated gene promoters in K*dm2a-CXXC*^*fl/fl*^ (left) or K*dm2a-CXXC*^*fl/fl*^ (right) mESCs following tamoxifen treatment. No promoters significantly changed in accessibility (p-adj < 0.05 and > 1.4-fold). (B) An illustration of the pausing index, the ratio of the average read density of RNAPII-NTD at the promoter and the average read density of RNAPII in the gene body. (C) Scatter plots comparing the log2 fold change in gene expression (cnRNA-seq) with the log2 fold change in RNAPII occupancy (ChIP-seq) following tamoxifen treatment of *Kdm2a/b-CXXC*^*fl/fl*^ mESCs, at CGI-associated gene promoters (left) or gene bodies (right). The solid line shows the linear regression, and the coefficient of determination (R^2^) and Spearman correlation coefficient (Cor) are annotated. (D) Boxplots showing the fold change in RNAPII occupancy at CGI-associated gene promoters (left) or gene bodies (right) following tamoxifen treatment of *Kdm2a/b-CXXC*^*fl/fl*^ mESCs, for all CGI genes or for genes that significantly increased in expression separated into quartiles according to their log2 fold change in expression (Q1 < Q2 < Q3 < Q4). (E) A metaplot showing RNAPII enrichment before (UNT) and after tamoxifen treatment (OHT) of *Kdm2a/b-CXXC*^*fl/fl*^ mESCs, for the top quartile of significantly upregulated genes (Q4 – as in (D)). (F) Heatmap analyses of the fold change in RNAPII, Ser5-RNAPII or Ser2P-RNAPII ChIP-seq signal following tamoxifen treatment of K*dm2a/b-CXXC*^*fl/fl*^ mESCs, for CGI-associated (n=14106) and non-CGI-associated (n=6527) gene promoters.

